# RIPPLE: learning nucleotide relations in an RNA foundation model

**DOI:** 10.64898/2026.11.23.768763

**Authors:** Yi Hao, Xudong Guo, Zixu Ran, Yannan Yang, Chun-Xia Zhao, Lachlan JM Coin, Christopher J Sweeney, Fuyi Li

## Abstract

RNA function depends on interactions between nucleotides that can be widely separated in sequence, yet capturing these relationships remains challenging for RNA language models. Existing approaches primarily learn nucleotide relationships implicitly from individual nucleotide reconstruction or rely on external structural information to model nucleotide interactions. Here we introduce RIPPLE, an RNA foundation model that learns to identify and represent sparse, predictively useful nucleotide relationships directly from sequence. Pretrained on 32.4 million RNA sequences, RIPPLE uses pairwise predictive utility to guide relational learning, transferring training-time supervision into a sequence-only model without requiring structural inputs at inference. RIPPLE improves overall performance across five downstream tasks spanning RNA family classification, ribosome loading prediction, secondary-structure prediction, three-dimensional contact prediction, and splice-site recognition. Mechanistic analyses further show that utility-guided learning improves the recovery of informative interactions and enhances the transferability of learned RNA representations.

## Introduction

RNA molecules encode biological activity not only through their nucleotide composition, but also through interactions between positions that can be widely separated along the primary sequence. Such interactions organise secondary and tertiary structure, alter accessibility and stability, and shape processes ranging from translation to splicing.^1-4^ A useful RNA representation should therefore capture both the local sequence context of individual nucleotides and the relational context through which distant positions constrain one another. Yet structural annotations remain sparse compared with the rapidly expanding volume of RNA sequence data,^5,6^ making self-supervised learning an attractive route to representations that can be reused across diverse prediction problems.

RNA language models have begun to exploit this asymmetry by pretraining Transformers on millions of unlabelled sequences.^7-16^ Sequence-centred models learn transferable representations from masked-token objectives,^7,8,12-16^ whereas more specialised approaches introduce motif-aware corruption, RNA-type conditioning, multiple-sequence alignments or predicted structural features.^7,9-11^ Collectively, these studies establish that large-scale pretraining can support RNA function and structure prediction, while systematic comparisons show that pretraining data, context length, tokenisation and downstream adaptation can materially affect model rankings.^17^ They also expose a central design choice. Models that operate only on sequence are broadly deployable, but conventional masked language modelling supervises the recovery of individual nucleotides and does not explicitly identify which pairwise relations are useful for that recovery. Conversely, models supplied with alignments or predicted structures can inject strong relational priors, but their inference path inherits the availability, computational cost, and errors of those external inputs.

Recent RNA models have incorporated dense pairwise attention biases, predicted folding features, and explicit geometric reasoning for structure prediction.^6,9,10,18^ These approaches demonstrate the value of relational information, but they leave open whether pairwise inductive bias can be internalised during pretraining and then retrieved directly from a single sequence. This question is important for two reasons. First, the number of possible nucleotide pairs grows quadratically with sequence length, whereas the interactions most useful for a particular prediction may be sparse. Second, a structural proposal is not the same as a universally useful relation: a candidate pair can help to predict the two nucleotides jointly, have little effect, or even be misleading in the model’s current representational state. A training signal should therefore measure pair-specific predictive utility rather than treating a proposed contact as an unquestioned label.

Here we introduce RIPPLE (**R**NA **I**nteraction-aware **P**airwise **P**retrained **L**anguage **E**ncoder), a sequence encoder designed to learn sparse relational computation while retaining sequence-only inference. RIPPLE augments the upper layers of a 26-layer Transformer with two coupled components. A Native Retriever selects a small set of candidate partners for each nucleotide from hidden states alone, and persistent pair states carry information along these sparse directed edges across layers. During pretraining, a separate objective branch constructs training-only pair candidates and estimates how much modelling a pair jointly improves the likelihood of its masked endpoints. This likelihood improvement, which we call Gain, distils a signed utility ranking into the Native Retriever. The objective branch can use folding-based proposals to diversify training examples, but it neither supplies hard structural labels nor participates in inference.

The resulting separation is deliberate: relational priors may guide the learning problem, but deployment requires only an RNA sequence. Pair-to-token feedback is bounded at every routed layer, and the total candidate mass assigned to sparse partners is capped, providing an explicit control on how much the relational pathway can perturb the sequence backbone. We designed controlled frozen-encoder evaluations spanning ncRNA family classification, mean ribosome load prediction, secondary-structure prediction, three-dimensional contact prediction, and multispecies splice-site recognition. These tasks probe wholesequence and pairwise representations under a common sequence-only inference contract.

## Results

### RIPPLE internalises relational supervision for sequence-only inference

RIPPLE was designed to learn sparse nucleotide relations during pretraining while preserving a sequence-only inference path (**Fig. 1**). The model comprises a 26-layer Transformer encoder, with its final eight layers augmented by sparse pair-aware modules that integrate relational information into nucleotide representations. At the interface between these stages, the Native Retriever selected up to eight candidate partners for every eligible nucleotide from the sequence representation alone. The choice of eight candidates reflects a balance between computational budget and the recall of true interaction partners. Each directed edge carried a 128-dimensional pair state that persisted across the pair-aware layers. Gated sparse attention then returned a bounded update to the sequence stream. The model therefore maintained an explicit relational memory without replacing the token representation or constructing a dense quadratic pair graph.

**Fig. 1:**
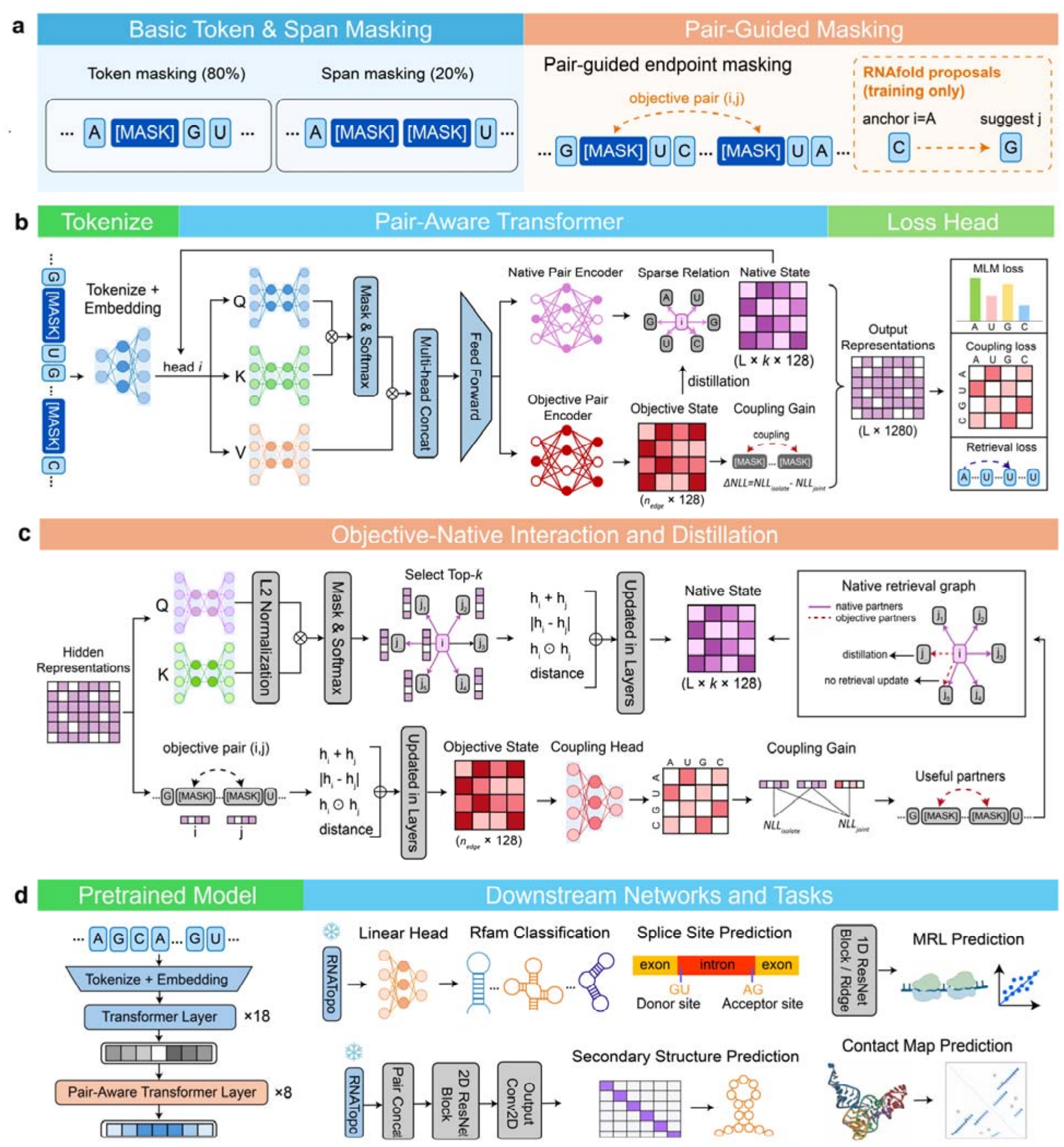
Architecture and training framework of RIPPLE. **a**, Conventional token and span masking is combined with pair-guided endpoint masking. RNAfold proposals diversify objective pairs during pretraining only and are not treated as structural labels. **b**, The 26-layer encoder comprises 18 conventional sequence layers followed by eight pair-aware layers. Native and objective pair encoders share the persistent pair-state machinery, whereas masked-language-modelling, coupling, and retrieval losses provide complementary supervision. **c**, For each objective edge, coupling gain is the reduction in endpoint negative log-likelihood under fixed-marginal joint rather than independent prediction. Detached gain rankings distil useful relations into the sequence-derived Native Retriever; objective edges already selected by the native path act as score references and are not updated by the retrieval loss. **d**, At deployment, the objective branch and folding proposals are removed. Frozen or task-adapted RIPPLE representations support sequence-level and relation-level downstream tasks from nucleotide sequence alone.

A separate objective branch supplied supervision for deciding which relations were useful. Pair-guided masking presented candidate endpoint pairs, and a fixed-marginal coupling head compared joint prediction of the two masked nucleotides with independent prediction. We defined the reduction in negative log-likelihood as Pair Gain (shown as coupling gain in **Fig. 1**). Preserving both single-nucleotide marginals prevented the coupling head from improving the score by changing either endpoint prediction in isolation. Detached Pair Gain rankings, with gradients stopped through the Gain values, instead trained the Native Retriever to favour candidate partners that improved joint prediction. Pair Gain therefore acted only as a teacher signal for retrieval and was not optimized through the retriever loss. The objective branch could receive folding-based proposals during pretraining, but these proposals were neither hard structural labels nor inputs to the deployed model.

RIPPLE was pretrained on 32.4 million RNA sequences and retained only the native sequence pathway at deployment. This separation yielded two empirically distinguishable internal readouts. If Pair Gain encoded endpoint compatibility, it should respond to nucleotide substitutions at fixed positions. By contrast, the Native Retriever ranked candidate partners directly from sequence-derived hidden representations, providing a relational neighbourhood in which broader organisation, such as continuity along helical regions, could be examined. The subsequent analyses tested these predictions independently before examining how both signals co-localised in individual RNAs.

### RIPPLE representations transfer across diverse RNA prediction tasks

We systematically evaluated RIPPLE representations’ transferability against 11 existing RNA language models across five downstream task families: ncRNA family classification, mean ribosome loading (MRL) prediction, secondary-structure prediction, three-dimensional contact prediction, and multispecies splice-site recognition (**Table 1**). To ensure controlled comparisons, all pretrained encoders were frozen and evaluated using matched task-specific prediction heads, with only sequence-derived nucleotide representations provided as downstream inputs. Dataset sources, processed splits and eligibility are summarized in **Supplementary Table 1**, and complete model-by-model results are reported in **Supplementary Tables 2-7**.

**Table 1:** Performance of RNA representations across five downstream task families. Values are macro-F1 for ncRNA classification, Spearman correlation and R^2^ for MRL, and exact F1-score for TS0, bpRNA-new, ArchiveII and RIVAS. Three-dimensional contacts use long-range P@L, and donor and acceptor splice-site predictions use F1. Best scores are bold; second-best scores are underlined (rankings based on unrounded values).

| Model | ncRNA | MRL |  | Secondary structure |  |  |  | 3D contacts |  | Splice-site |  |
|---|---|---|---|---|---|---|---|---|---|---|---|
|  | Macro-F1 | Ridge<br>Spearman | ResNet<br>R<sup>2</sup> | TS0<br>F1 | bpRNA-new<br>F1 | ArchiveII<br>F1 | RIVAS<br>F1 | Strong<br>P@L | Probe<br>P@L | Donor<br>F1 | Acceptor<br>F1 |
| RNABERT | 0.533 | 0.359 | 0.651 | 0.404 | 0.399 | 0.272 | 0.369 | 0.389 | 0.204 | 0.551 | 0.549 |
| RNA-FM | 0.840 | 0.423 | 0.659 | 0.579 | 0.436 | 0.290 | 0.506 | 0.518 | 0.364 | 0.660 | 0.669 |
| RNA-MSM | 0.532 | 0.333 | 0.603 | 0.442 | 0.410 | 0.261 | 0.348 | 0.396 | 0.275 | 0.631 | 0.725 |
| Uni-RNA | 0.573 | 0.466 | 0.613 | 0.497 | 0.470 | 0.353 | 0.424 | 0.459 | 0.321 | 0.662 | 0.692 |
| RiNALMo | 0.851 | <u>0.503</u> | 0.557 | 0.595 | 0.459 | <u>0.590</u> | 0.588 | 0.558 | <u>0.423</u> | <u>0.694</u> | 0.715 |
| ERNIE-RNA | <u>0.906</u> | 0.418 | 0.656 | <u>0.660</u> | **0.559** | 0.576 | <u>0.655</u> | <u>0.565</u> | 0.334 | 0.682 | 0.701 |
| RNAErnie | 0.720 | 0.436 | 0.594 | 0.502 | 0.413 | 0.395 | 0.408 | 0.446 | 0.325 | 0.660 | 0.545 |
| BiRNA-BERT | 0.648 | 0.423 | <u>0.666</u> | 0.490 | 0.456 | 0.299 | 0.442 | 0.458 | 0.191 | 0.647 | 0.677 |
| HydraRNA | 0.632 | **0.561** | 0.510 | 0.521 | 0.479 | 0.364 | 0.433 | 0.517 | **0.460** | 0.681 | **0.765** |
| NucleicBERT | 0.565 | 0.385 | 0.597 | 0.566 | 0.439 | 0.458 | 0.474 | 0.355 | 0.286 | 0.670 | <u>0.754</u> |
| RNAret | 0.595 | 0.384 | 0.472 | 0.394 | 0.394 | 0.334 | 0.348 | 0.379 | 0.296 | 0.652 | 0.645 |
| RIPPLE | **0.925** | 0.485 | **0.689** | **0.671** | <u>0.509</u> | **0.599** | **0.676** | **0.575** | 0.422 | **0.710** | 0.736 |

RIPPLE ranked first in seven of the eleven primary evaluation metrics, second in one and third in three. For ncRNA family classification, RIPPLE achieved the highest macro-F1 of 0.925 (accuracy, 0.974). In MRL prediction, RIPPLE achieved the highest R^2^ of 0.689 with a nonlinear residual-network decoder and the third-highest Spearman correlation of 0.485 with ridge regression. For multispecies splice-site recognition, RIPPLE achieved the highest donor F1 of 0.710 and the third-highest acceptor F1 of 0.736.

The advantages of RIPPLE were particularly evident in structure-related tasks. Across four secondary-structure benchmarks, RIPPLE achieved exact F1 scores of 0.671, 0.509, 0.599 and 0.676 on bpRNA-TS0, bpRNA-new, ArchiveII and RIVAS, respectively, ranking first on three benchmarks and second on bpRNA-new. Its performance on bpRNA-new and the family-held-out ArchiveII evaluation further supported the transferability of its representations across distinct structural datasets. In three-dimensional contact prediction, RIPPLE achieved the highest long-range P@L of 0.575 using a nonlinear decoder and the third-highest P@L of 0.422 with a lightweight probe. These results indicate that information relevant to nucleotide-pair relationships was accessible from frozen sequence representations without directly exposing learned pair states or retrieval scores to downstream predictors.

Together, these findings demonstrate the broad transferability of RIPPLE representations across RNA prediction tasks, with particularly strong performance in secondary-structure prediction. This motivated further investigation into the relational information learned during pretraining and its correspondence with biologically meaningful RNA interactions.

### Frozen RIPPLE representations organise ncRNA family space

We first tested whether the frozen encoder retained whole-sequence organisation beyond the pairwise training objective. A two-dimensional projection of mean-pooled RIPPLE embeddings for all 18,347 test RNAs in the 12 most frequent ncRNA families formed multiple family-enriched neighbourhoods (**Fig. 2a,b**). The same organisation was recovered without relying on the two-dimensional projection. In a balanced evaluation set containing up to 400 sequences per family (n = 4,766), ten-nearest-neighbour family purity in the high-dimensional embedding space was 0.990 overall. Per-family purity ranged from 0.967 to 1.000 (**Fig. 2c**), showing that local family coherence was not confined to the largest classes. Family-resolved test counts and high-dimensional neighbourhood purity are reported in **Supplementary Table 8**.

**Fig. 2:**
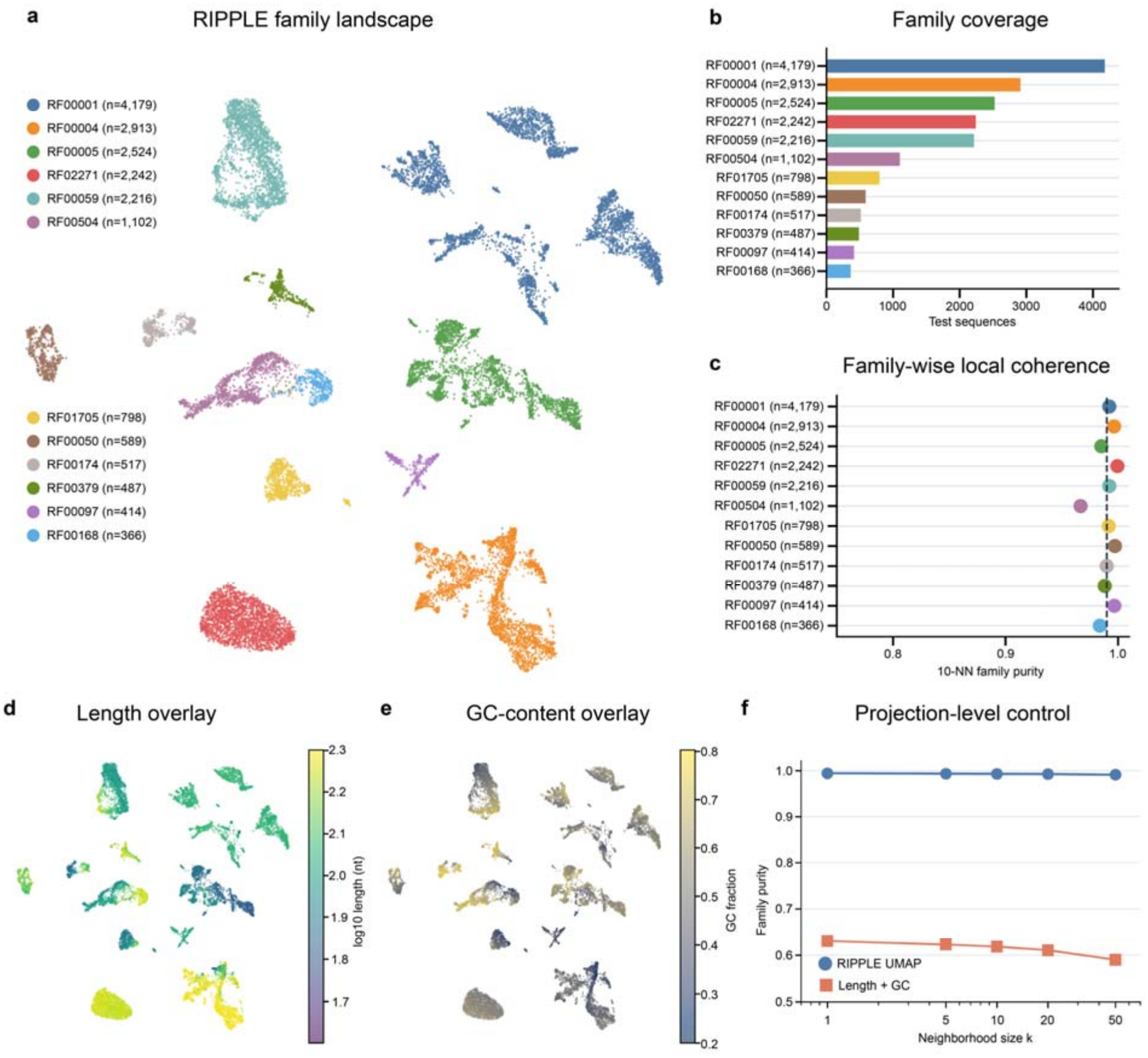
Frozen RIPPLE embeddings form family-enriched neighbourhoods. **a**, Uniform manifold approximation and projection (UMAP) of mean-pooled frozen RIPPLE embeddings after principal-component reduction to 64 dimensions for all 18,347 test RNAs in the 12 most frequent ncRNA families. Each point is one RNA, and colours denote Rfam family labels. **b**, Number of test sequences contributed by the 12 families. **c**, Familywise ten-nearest-neighbour purity calculated in the high-dimensional embedding space on a balanced subset containing up to 400 sequences per family (n = 4,766). The dashed line denotes the mean across families. **d,e**, The same UMAP coordinates coloured by log10 sequence length (**d**) or GC fraction (**e**). **f**, Family purity in the two-dimensional RIPPLE projection and in a two-dimensional length-plus-GC control across neighbourhood sizes k = 1, 5, 10, 20 and 50 (balanced n = 4,766).

Sequence length and GC content varied within most projected family regions and formed gradients that crossed local cluster boundaries (**Fig. 2d,e**). These coarse features therefore contributed to the geometry but did not reproduce the family labels. In the balanced subset, family purity in the RIPPLE projection decreased only from 0.995 at k = 1 to 0.991 at k = 50. A two-dimensional control built from sequence length and GC content decreased from 0.630 to 0.590 over the same neighbourhood sizes (**Fig. 2f**). The separation persisted as the neighbourhood expanded, which argued against a small number of isolated points driving the effect.

Agreement between the high-dimensional analysis and the projection-level control addressed two common ambiguities in RNA embedding maps, visual separation introduced by the projection algorithm and clustering dominated by gross sequence composition. The results established that the frozen representation organised RNAs by family-associated sequence regularities beyond these two features.

### Pair Gain reveals biologically organised pairwise utility

We next tested whether Pair Gain separated annotated nucleotide pairs from controls that preserved simpler properties of the evaluated positions. Across a deduplicated union of four secondary-structure sources, comprising 8,041 RNAs and 146,698 sampled edges, annotated pairs had an RNA-level mean Pair Gain of 0.697. Distance-matched false pairs and random pairs had mean values of 0.231 and -0.067, respectively (**Fig. 3a**). The annotated-pair advantage over distance-matched controls was 0.465 (95% confidence interval (CI), 0.458-0.472). At the edge level, Pair Gain distinguished annotated pairs from all controls with an area under the precision-recall curve (AUPRC) of 0.753 and from distance-matched controls with an AUPRC of 0.795. Pair Gain retained discriminatory power against distance-matched false pairs, indicating that its signal was not solely explained by sequence separation.

**Fig. 3:**
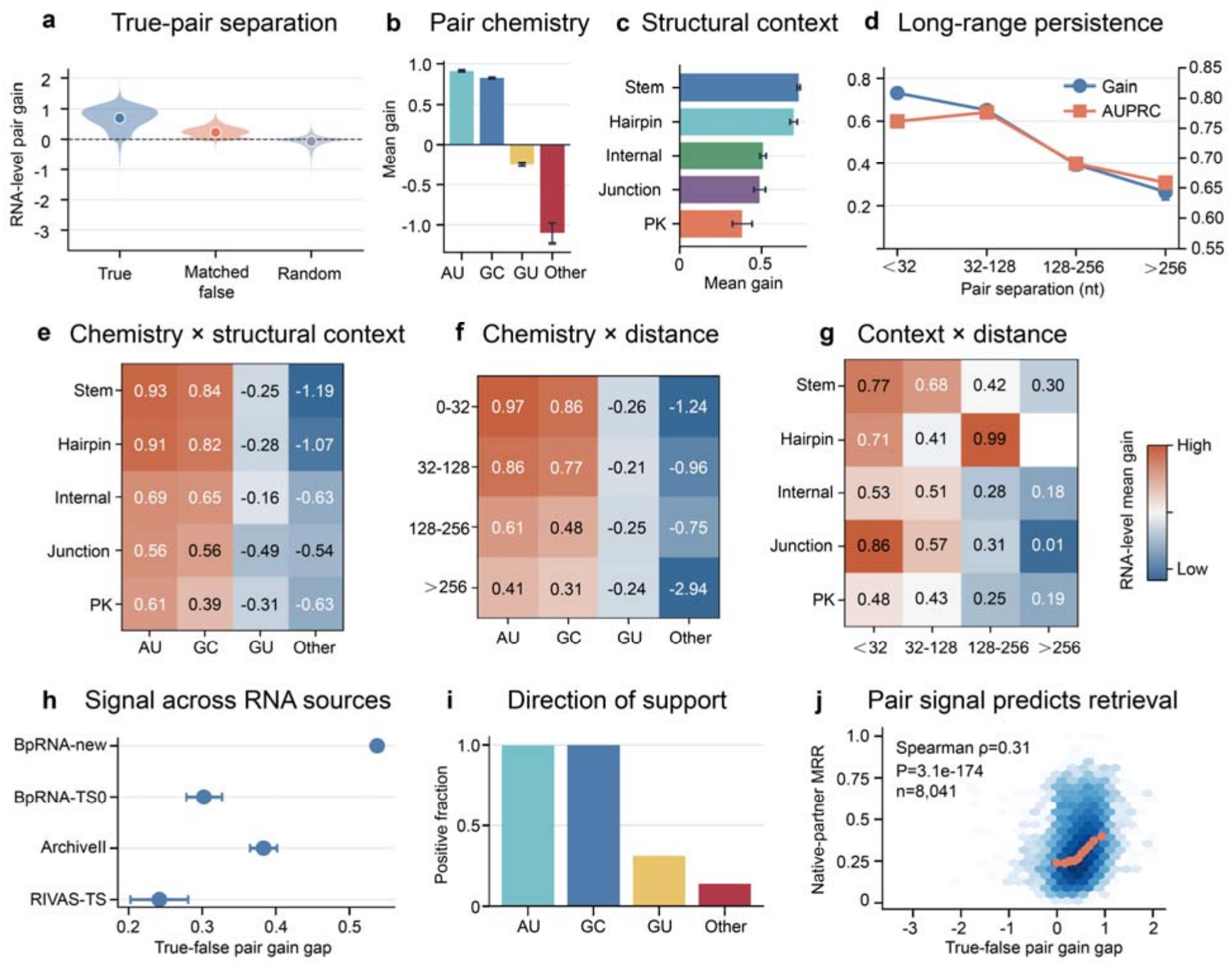
Pair Gain captures chemistry, context and range of RNA pairing. **a**, Distributions of per-RNA mean Pair Gain for annotated pairs, distance-matched false candidates and random candidates across a deduplicated union of four secondary-structure sources (n = 8,041 RNAs per group; 146,698 sampled edges). White circles denote medians and thick and thin lines denote interquartile ranges and observed ranges, respectively. **b**, RNA-level mean Pair Gain for annotated AU, GC, GU and other pairs (n = 6,806, 7,834, 3,582 and 623 RNAs). **c**, RNA-level mean Pair Gain in stem, hairpin, internal-loop, junction and pseudoknot contexts (n = 8,026, 3,715, 3,830, 1,451 and 443 RNAs). **d**, Mean annotated-pair Gain and edge-level AUPRC across sequence-separation bins ( 32, 32-128, 128-256 and 256 nucleotides; n = 7,906, 6,863, 1,176 and 595 RNAs). **e-g**, RNA-level mean Pair Gain jointly stratified by pair chemistry and structural context (**e**), pair chemistry and separation (**f**), or structural context and separation (**g**); heatmaps share a zero-centred colour scale. **h**, RNA-level annotated-minus-matched Pair Gain by source (n = 5,235 BpRNA-New, 1,173 BpRNA-TS0, 1,249 ArchiveII and 384 RIVAS-TS RNAs). **i**, Fraction of annotated edges with positive Pair Gain by pair chemistry (n = 16,013 AU, 27,027 GC, 4,964 GU and 980 other edges). **j**, Association between the RNA-level annotated-minus-matched Pair Gain gap and native-partner mean reciprocal rank (n = 8,041 RNAs). Hexagons show RNA density, and the orange line connects medians across Pair Gain deciles. Error bars in **b-d** and **h** denote 95% confidence intervals from 2,500 RNA-level bootstrap resamples. Pair Gain is measured in nats.

Pair Gain was strongly organised by nucleotide chemistry. RNA-level mean values were 0.911 (95% CI, 0.900-0.923) for AU pairs and 0.824 (95% CI, 0.815-0.833) for GC pairs. GU pairs and other annotated states instead had mean values of -0.249 (95% CI, -0.267 to -0.230) and -1.101 (95% CI, -1.227 to - 0.977), respectively (**Fig. 3b**). The sign of the edge-level effect showed the same ordering: Pair Gain was positive for 99.8% of AU and 100.0% of GC edges, compared with 31.2% of GU and 13.9% of other edges (**Fig. 3i**). Pair Gain thus represented a signed utility score rather than an unsigned measure of proximity or annotation status. Pair-chemistry statistics and positive-Gain fractions are detailed in **Supplementary Table 9**.

Structural context and sequence range further stratified this chemistry-dependent signal. Mean Pair Gain was 0.727 in stems and 0.696 in hairpins, compared with 0.508 in internal loops, 0.487 in junctions and 0.381 in pseudoknots (**Fig. 3c**). Chemistry-by-context maps retained positive AU and GC values in every context, whereas GU and non-canonical states remained weaker or negative (**Fig. 3e**). Mean Pair Gain decreased from 0.733 (95% CI, 0.724-0.743) below 32 nucleotides to 0.264 (95% CI, 0.227-0.301) beyond 256 nucleotides, and AUPRC decreased from 0.762 to 0.659 (**Fig. 3d**). Canonical pairs nevertheless retained positive values in every distance stratum (**Fig. 3f,g**), showing that the coupling signal extended beyond local sequence neighbourhoods. Context- and distance-stratified values are reported in **Supplementary Tables 10-11**.

The annotated-versus-matched Pair Gain difference was positive in all four data sources (**Fig. 3h**), arguing against a single collection as the origin of the aggregate effect. At the RNA level, this difference correlated with native partner mean reciprocal rank (Spearman’s ρ = 0.306, P = 3.08×10^-174^, n = 8,041; **Fig. 3j**). The association was consistent but moderate. Pair Gain and native retrieval were therefore linked by training, yet they were not interchangeable measurements: the coupling objective quantified the utility of a tested relation, whereas the retriever had to rank candidate partners from the sequence representation before the pair was evaluated. Source-specific contrasts and Gain-gap decile summaries are given in **Supplementary Tables 12-13**.

### Counterfactual and compensatory perturbations recover nucleotide compatibility

We then tested whether Pair Gain responded to endpoint identity while positions and surrounding sequence context were held fixed. From each of 6,445 eligible RNAs, selected independently of model output, we chose the longest-range Watson-Crick pair supported by at least one adjacent stacked pair. All 16 ordered nucleotide states were evaluated at the same two positions. The four Watson-Crick states had median Pair Gain values of 1.036-1.060, whereas the two GU states had medians of -0.133 and -0.205. The ten mismatch states ranged from -0.572 to -2.281 (**Fig. 4a**). Within RNAs, the median Watson-Crickminus-mismatch contrast was 2.895 (95% CI, 2.854-2.938), directly linking the sign and magnitude of Pair Gain to endpoint compatibility. The complete 16-state Pair Gain summary is provided in **Supplementary Table 14**.

**Fig. 4:**
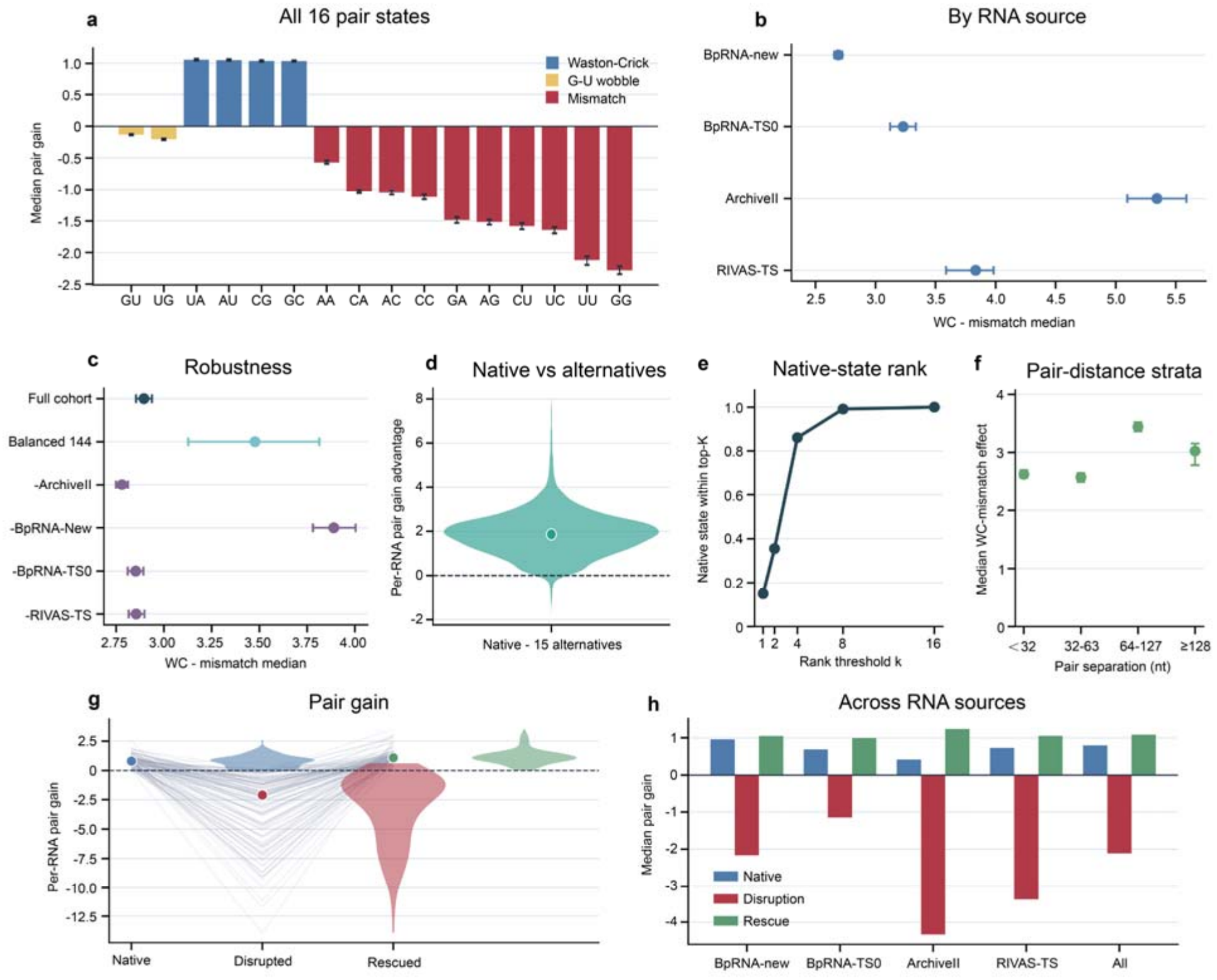
Counterfactual and compensatory perturbations expose pair compatibility. **a**, Median Pair Gain for all 16 ordered nucleotide states evaluated at the same masked positions in 6,445 RNAs. Colours distinguish Watson-Crick states, GU wobble states and mismatches. For each RNA, the target was the longest-range Watson-Crick pair supported by at least one adjacent stacked pair. **b**, Within-RNA contrast between the median Pair Gain of the four Watson-Crick states and that of the ten mismatch states by source (n = 586 ArchiveII, 4,425 BpRNA-New, 1,058 BpRNA-TS0 and 376 RIVAS-TS RNAs); GU states are excluded. **c**, The same contrast in the complete cohort, a source-balanced sensitivity set of 144 RNAs (36 per source) and four leave-one-source-out cohorts. **d**, Per-RNA advantage of the native state over the median of the other 15 states. **e**, Cumulative fraction of RNAs for which the native state ranks within the top k of the 16 states. f, Watson-Crick-minus-mismatch contrast across pair-separatio strata (<32, 32-63, 64-127 and =128 nt; n = 1,021, 2,369, 2,714 and 341 RNAs, respectively). **g**, Pair Gain for native, single-disruption and compensatory-rescue variants in the balanced 144-RNA set. Pale lines connect variants of the same RNA; violins show distributions and coloured points indicate medians. **h**, Median Pair Gain for the three variants by source and across all sources. Error bars, where shown, denote 95% confidence intervals from 4,000 RNA-level bootstrap resamples.

The compatibility contrast was robust to source composition and sequence range. Source-specific medians ranged from 2.690 in BpRNA-New to 5.343 in ArchiveII, and the effect remained positive when each source was excluded in turn (**Fig. 4b,c**). A balanced sensitivity set containing 36 RNAs from each source produced a median contrast of 3.477 (95% CI, 3.127-3.814). The contrast also remained positive in every separation band, including pairs separated by at least 128 nucleotides (median, 3.023; 95% CI, 2.780-3.150; **Fig. 4f**). These controls showed that the state ordering was not created by the dominant source or by short-range pairs.

The native nucleotide state exceeded the median of the other 15 states by 1.861 Pair Gain units and showed a positive advantage in 99.1% of RNAs (**Fig. 4d**). Native states ranked within the top four alternatives in 86.1% of RNAs and within the top eight in 99.1% (**Fig. 4e**). Their top one frequency was only 15.0%. The objective therefore recovered a compatibility class more consistently than a unique ordered state, which is expected when several Watson-Crick substitutions remain plausible at the same positions.

A paired sequence intervention tested whether the effect was reversible. For one canonical stacked pair in each of 144 source-balanced RNAs, a single substitution reduced median Pair Gain from 0.801 to -2.111, whereas a second substitution that restored complementarity increased it to 1.086 (**Fig. 4g**). The median rescue-minus-disruption effect was 3.471 (95% CI, 2.885-3.972), and rescue increased Pair Gain in 99.3% of RNAs. Source-specific rescue fractions ranged from 97.2% to 100.0%(**Fig. 4h**). In contrast, the Native Retriever probability assigned to the target coordinate showed no corresponding rescue effect (median difference, 2.5 × 10□□; 95% CI, -3.0 × 10□□ to 7.3 × 10□□). The perturbation therefore changed the chemistry-sensitive coupling score without producing the same response in the context-sensitive retriever. These in silico interventions establish a model-internal compatibility mechanism, rather than physical rescue of an experimentally measured RNA structure. Source-stratified compatibility contrasts and rescue statistics are provided in **Supplementary Table 15**.

### The Native Retriever resolves helix topology from sequence

We evaluated the Native Retriever at the level of helix organisation rather than endpoint chemistry. The analysis included every RNA between 60 and 240 nucleotides with at least four annotated pairs, giving 6,417 RNAs, 158,386 pairs and 36,771 stems across four sources. Reference base-pair annotations were used only after sequence-only inference to define evaluation strata and were never provided to the Native Retriever as input. RNA-macro recall of the annotated partner increased from 0.289 at rank 1 to 0.532 at rank 4, 0.671 at rank 8 (Recall@8) and 0.806 at rank 16 (**Fig. 5a**). Recall@8 ranged from 0.552 on RIVAS-TS to 0.778 on ArchiveII (**Fig. 5e**). The retrieval signal was therefore consistent across collections despite differences in annotation source and RNA composition.

**Fig. 5:**
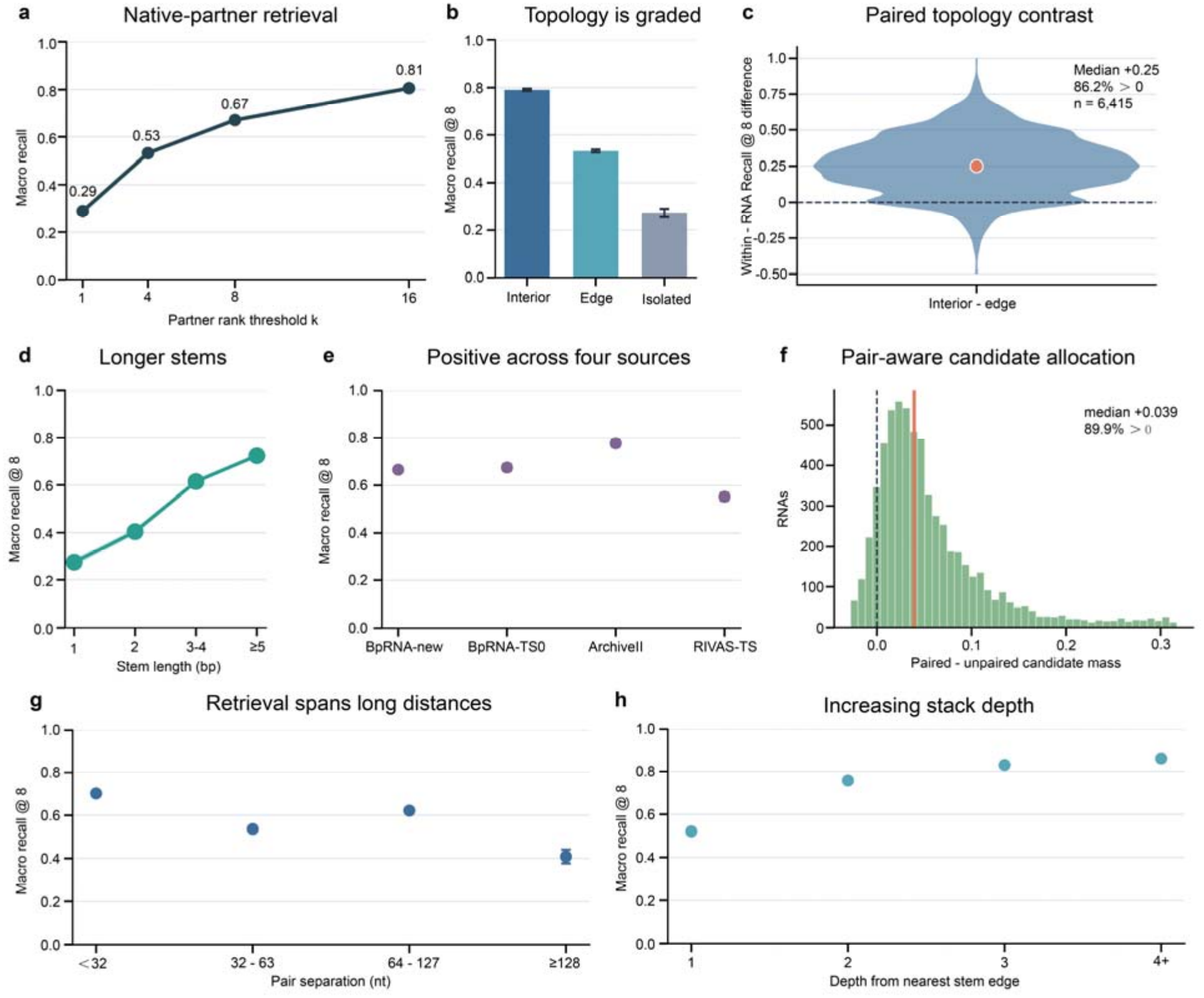
The sequence-only Native Retriever encodes helix topology. **a**, RNA-macro recall of annotated partners as a function of retrieval depth, referred to as Recall@X for X =1,4,8,16 (n = 6,417 RNAs). Recall is calculated per RNA before averaging. **b**, Recall@8 for pairs in stem interiors, at stem edges or outside multi-pair stems (n = 6,415, 6,417 and 2,379 RNAs). **c**, Within-RNA interior-minus-edge difference in Recall@8 for RNAs containing bot classes (n = 6,415). The coloured point denotes the median. **d**, Recall@8 across stem-length bins of 1, 2, 3-4 and at least 5 base pairs (n = 2,379, 2,835, 5,375 and 6,102 RNAs). **e**, Recall@8 across BpRNA-New, BpRNA-TS0, Ar-chiveII and RIVAS-TS (n = 4,425, 1,030, 586 and 376 RNAs). **f**, Distribution of the within-RNA difference in mean Native Retriever candidate mass between paired and unpaired anchors (n = 6,417 RNAs). The red line denotes the median and the dashed line denotes zero. **g**, Recall@8 across pair-separation bins of 32, 32-63, 64-127 and 128 nucleotides (n = 6,316, 5,172, 3,248 and 417 RNAs). **h**, Recall@8 by stack depth from the nearest stem edge (n = 6,417, 6,415, 6,102 and 4,140 RNAs for depths 1, 2, 3 and 4). Point and bar estimates in a,b,d,e,g, and h show means with 95% confidence intervals from 4,000 RNA-level bootstrap resamples. The analysis comprises 158,386 annotated pairs in 36,771 stems from RNAs of 60-240 nucleotides containing at least four annotated pairs. Structural annotations were used only after inference to define evaluation strata.

Recovery was strongly graded by a pair’s position within a helix. Mean Recall@8 was 0.791 (95% CI, 0.786-0.796) for stem-interior pairs, 0.533 (95% CI, 0.527-0.539) for stem-edge pairs, and 0.273 (95% CI, 0.256-0.290) for isolated pairs (**Fig. 5b**). This ordering was also present within individual molecules. Among 6,415 RNAs containing both interior and edge positions, the median interior-minus-edge difference was 0.250, and 86.2% showed a positive difference (**Fig. 5c**). Requiring reciprocal recovery strengthened the same hierarchy: bidirectional Recall@8 was 0.598 for interior pairs, 0.323 for edge pairs and 0.129 for isolated pairs. The gradient therefore could not be attributed only to different RNAs contributing to each category.

Two independent measures of helix support produced concordant trends. Recall@8 increased from 0.273 for isolated pairs to 0.724 for stems containing at least five pairs (**Fig. 5d**). It also increased from 0.521 at the first stack layer to 0.861 at depth four or greater (**Fig. 5h**). Stem length measures the extent of the complete helix, whereas stack depth measures how far a pair lies from the nearest stem edge, with an edge pair assigned depth 1 and each additional consecutive base-pair step increasing the depth by one. Their agreement showed that native retrieval was strongest for relations embedded within repeated, internally supported pairing patterns.

The Native Retriever also allocated more of its finite candidate mass to paired than to unpaired anchors. The within-RNA paired-minus-unpaired difference had a median of 0.039 (95% CI, 0.038-0.041) and was positive in 89.9% of RNAs (**Fig. 5f**). Sequence distance constrained, but did not determine, retrieval. Mean Recall@8 was 0.702 below 32 nucleotides, 0.537 at 32-63 nucleotides, 0.622 at 64-127 nucleotides and 0.409 at 128 nucleotides or more (**Fig. 5g**). The non-monotonic middle bins and the strong effects of stem position and stack depth showed that the retriever was not simply ranking the nearest eligible nucleotide. Together, these results identify it as a topology-sensitive candidate selector rather than a calibrated base-pair probability or a direct readout of endpoint compatibility. Source- and helix-topology-stratified native recall results are detailed in **Supplementary Table 16**.

### Learned relational signals localise within individual RNAs

We next examined how Pair Gain and native retrieval co-localised within individual molecules (**Fig. 6**). Three RNAs were selected by deterministic criteria to cover long-range, mixed-chemistry and compactstem regimes. The 117-nucleotide bpRNA_RFAM_6139 contained 21 annotated pairs, all separated by at least 64 nucleotides, and showed an annotated-versus-matched Pair Gain gap of 1.902 and native-partner Recall@8 of 0.952. The 81-nucleotide compact stem network bpRNA_RFAM_21434 showed a Gain gap of 1.108 and Recall@8 of 0.963. The 88-nucleotide mixed-pair helix URS0000D69221_12908_1 showed a smaller Gain gap of 0.925 and Recall@8 of 0.632 (**Fig. 6a-c,g**). The three case-level readouts are summarised in **Supplementary Table 17**.

**Fig. 6:**
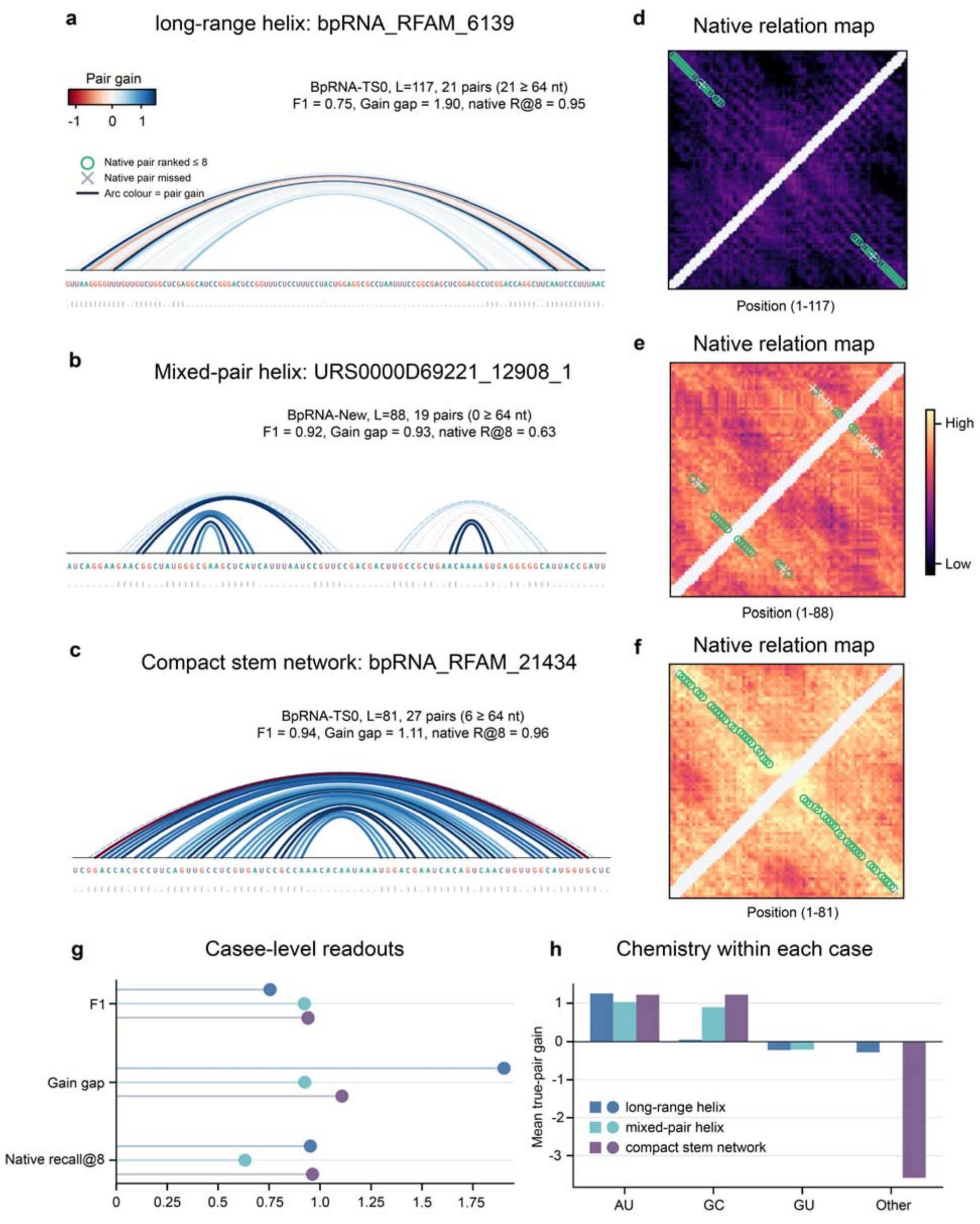
Real RNA examples connect pair compatibility to local secondary structure. **a-c**, Annotated secondary-structure arcs for the long-range helix bpRNA_RFAM_6139 from BpRNA-TS0 (**a**; 117 nucleotides, 21 pairs), the mixed-pair helix URS0000D69221_12908_1 from BpRNA-New (**b**; 88 nucleotides, 19 pairs) and the compact stem network bpRNA_RFAM_21434 from BpRNA-TS0 (**c**; 81 nucleotides, 27 pairs). Arc colour encodes Pair Gain; green open circles and grey crosses denote annotated pairs whose native partner was recovered or missed, respectively, within the top eight sequence-derived candidates in at least one direction. Nucleotide identities and dot-bracket annotations are shown below each diagram. Gain gap is the RNA-level difference between mean Pair Gain for annotated and distance-matched false pairs. **d-f**, Sequence-only Native Retriever relation maps for the RNAs in **a-c**. Colours encode the base-10 logarithm of native-relation probability; green circles mark recovered annotated pairs, grey crosses mark missed pairs, and the invalid self-relation diagonal is masked. These maps visualise internal relation probabilities and are not independently decoded structures. **g**, Secondary-structure F1, Gain gap and native-partner Recall@8 for the three RNAs. **h**, Mean Pair Gain for annotated AU, GC, GU and other pairs in each RNA; absent bars indicate that a case contains no annotated pair in that class.

The case-level readouts separated relational recovery from final secondary-structure prediction. bpRNA_RFAM_6139 had high native-partner recovery but a lower F1 of 0.755, whereas the mixed-pair case reached an F1 of 0.923 despite recovering only 63.2% of annotated partners within the top eight candidates. The compact network combined high values for both measures (F1, 0.941; Recall@8, 0.963). Native retrieval and the downstream structure readout therefore captured related but non-identical stages of the model. A downstream decoder could recover a structure without every annotated edge appearing in the sparse candidate set, while accurate candidate recovery alone did not guarantee the highest final F1.

Arc diagrams and native relation maps localised the population-level mechanisms. Positive AU and GC Pair Gain followed coherent stem patterns in both compact and long-range helices, while retrieved edges traced the corresponding off-diagonal relation bands (**Fig. 6a-f**). Chemistry-resolved values exposed weaker non-canonical support. In the mixed-pair helix, mean annotated-pair Gain was 1.035 for AU, 0.895 for GC and -0.207 for GU pairs (**Fig. 6h**). The cases thus reproduced the division of labour established by the cohort analyses: Pair Gain varied with endpoint compatibility, whereas the Native Retriever outlined the surrounding helix configuration. Because the cases were selected for mechanistic coverage, they demonstrate localisation of the signals but do not estimate their prevalence across RNA families. Case-level chemistry-resolved Gain values are reported in **Supplementary Table 18**.

## Discussion

RNA language models must capture both nucleotide-level sequence information and interactions between positions that may be widely separated along the sequence. However, conventional masked language modelling does not explicitly distinguish which nucleotide relationships contribute to prediction. RIPPLE addresses this limitation by introducing pairwise predictive utility as a source of relational supervision. Rather than treating proposed nucleotide pairs as structural labels, the model evaluates their contribution to joint nucleotide prediction and transfers this information into a sequence-derived retrieval mechanism. By integrating sparse, persistent pair representations into a Transformer backbone, RIPPLE combines relational modelling with conventional sequence learning while retaining an inference pathway that requires no external structural inputs. This framing aligns with broader biomolecular and genomic language-model studies showing that self-supervised sequence pretraining can yield transferable structural and regulatory representations.^19-21^

The controlled downstream evaluations demonstrated the broad transferability of RIPPLE representations. Across five task families, RIPPLE ranked first in seven of eleven primary metrics, second in one and third in three. Its advantages were particularly evident in secondary-structure and three-dimensional contact prediction, where modelling nucleotide relationships is directly relevant to the prediction objectives. Competitive results in ncRNA family classification, ribosome loading prediction and multispecies splicesite recognition further suggested that incorporating relational learning did not compromise the representation of broader sequence-level information. Nevertheless, performance varied across datasets and decoder architectures, indicating that the benefits of relational pretraining are not uniform across all prediction settings. The result is also consistent with thermodynamic, probabilistic and deep-learning RNA folding methods, in which explicit pairwise representations are central to secondary-structure prediction.^22-25^ Long-context sequence models have likewise demonstrated that dispersed regulatory and splice determinants can be learned from primary sequence.^26,27^

Mechanistic analyses provided further insight into the relational information captured by RIPPLE. Pair Gain distinguished annotated nucleotide pairs from matched controls and showed strong sensitivity to nucleotide identity. Counterfactual substitutions and compensatory mutations further revealed that this signal reflected nucleotide compatibility, with restoration of complementarity recovering the predictive benefit of joint modelling. In contrast, the Native Retriever exhibited a stronger association with structural context, preferentially recovering relations within well-supported stems and stacked regions. At the RNA level, the annotated-minus-matched Pair Gain gap showed a moderate positive correlation with native-partner mean reciprocal rank (Spearman’s ρ = 0.306; **Fig. 3j**). This association, together with their different responses to nucleotide perturbations, suggested that the two signals captured complementary aspects of nucleotide relationships. These findings support a distinction between evaluating the predictive utility of a candidate relation and identifying informative partners from sequence representations alone. Evolutionary-coupling studies have independently shown that coordinated nucleotide variation can recover secondary, tertiary and functional contacts, although those approaches require homologous sequence alignments.^28,29^ The chemistry-dependent separation among Watson-Crick, GU and other states is also consistent with established geometric and biochemical distinctions among RNA base-pair classes.^30,31^

Several limitations also point to opportunities for further development. Pair Gain was designed as a predictive training signal rather than as a direct biophysical measurement, and its stronger responses to canonical Watson-Crick pairs than to GU and non-canonical interactions suggest that future work could incorporate broader relational supervision or complementary objectives that better capture non-canonical and context-dependent interactions. Native retrieval showed stronger recovery for pairs embedded within well-supported stems than for isolated or some long-range relations. This difference may partly reflect the greater contextual support available within helices, where neighbouring pairing patterns provide repeated relational evidence, whereas isolated and distant interactions are inherently less redundant and must be distinguished from a larger set of alternative partners. Future work could explore adaptive candidate allocation or multiscale relational representations to better accommodate such sparse and context-dependent relationships while retaining computational efficiency. Finally, RIPPLE currently uses folding-derived proposals as one source of relational prior during pretraining. These proposals serve to expose potentially informative nucleotide relationships to the training objective rather than constituting an inference-time requirement. The framework is therefore not inherently tied to a particular folding algorithm and could incorporate alternative structure predictors, experimentally derived relationships or other nucleotide-association priors. Potential training-time priors could include family-specific sequence-structure infor-mation from Rfam covariance models and Infernal^32,33^, or experimentally informed structural constraints derived from chemical probing approaches such as DMS-MaPseq.^34^ Future work could examine how the source, diversity and coverage of such relational priors affect the efficiency with which external relational knowledge is internalised into the sequence-only model. RIPPLE deliberately operates without epigenetic or cellular-context inputs, allowing relational information to be inferred from sequence alone. This portability is an advantage for foundation-model pretraining and deployment, and future extensions could incorporate such contextual signals when available to model how sequence-encoded relational preferences are modulated across cellular states.

Together, these findings suggest that learning the predictive utility of nucleotide relationships offers a promising direction for RNA foundation models. More broadly, RIPPLE illustrates how relational and structure-relevant information can be internalised during pretraining while preserving sequence-only inference, moving RNA foundation models beyond representations dominated by individual nucleotide context. Such models may provide a foundation for more predictive and ultimately design-oriented approaches to RNA biology, including the interpretation of structure-function relationships and the development of RNA engineering and therapeutic applications. Related advances in protein structure prediction further demonstrate the potential of machine learning to connect sequence information with molecular structure.^35,36^ Future work could extend this framework to longer RNA sequences, more diverse interaction types and less well-characterised RNA families, while examining how learned relational representations contribute to biological prediction and design under different adaptation settings.

## Methods

### Study design and sequence-only inference contract

RIPPLE was designed around the hypothesis that pair-specific supervision can be used during pretraining to internalise sparse relational representations without imposing an external structural pipeline at deployment. The model therefore contains two deliberately separated computational paths. The native path consists of the sequence Transformer, the Native Retriever, persistent sparse pair states and bounded pair-to-token feedback. This path is retained for all downstream evaluations. The objective path samples additional edges, estimates their pairwise predictive utility and supervises retrieval during pretraining; it is removed after pretraining.

At inference, RIPPLE consumes nucleotide tokens and padding information only. It does not require RNAfold output, base-pair probabilities, a multiple-sequence alignment, structural tokens or a teacher model. The implementation contains approximately 520 million trainable parameters, including about 512 million in the sequence backbone and 8.6 million in the pair-aware modules.

### RIPPLE architecture

#### Tokenisation and sequence backbone

Sequences were converted to RNA alphabet by replacing thymine with uracil. The vocabulary comprised nine tokens: <pad>, <cls>, <sep>, <mask>, A, U, C, G and N, in which the <cls> token marks the sequence-level classification and start position, whereas <sep> marks the end of the sequence. Characters outside A/U/C/G were mapped to N. Input sequences contained at most 1,024 biological nucleotides; the two boundary tokens gave a maximum model length of 1,026.

The backbone was a 26-layer Transformer encoder ^37^ with hidden width 1,280, 20 attention heads and feed-forward width 5,120. It used rotary positional embeddings,^38^ GELU activations, dropout of 0.1, atten-tion dropout of 0.1 and layer-normalization epsilon of 10^-12^. Linear and embedding weights were initial-ized with standard deviation 0.02. Each block used post-residual layer normalization. On compatible CUDA inputs in bfloat16 or float16, attention used a packed variable-length Flash Attention path.^39^ The first 18 layers (indices 0-17) were conventional sequence layers. The final eight layers (indices 18-25) were pair-aware.

#### Notation and conventions

Unless otherwise stated, *i* and *j* index nucleotide positions, *l* indexes pair-aware layers and m indexes sparse-attention heads. The vector *h*_*i*_ is the 1,280-dimensional sequence hidden state at position *i*, and 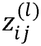is the 128-dimensional directed pair state from anchor *i* to candidate partner *j* after pair-aware layer *l. LN* denotes layer normalization; matrices denoted by *w* are learned linear projections; ‖*x* ‖2 is the Euclidean norm of vector *x*; a centred dot denotes an inner product; [*x*‖*y*] denotes concatenation; and ⊙ denotes element-wise multiplication.

#### Transformer sequence modelling

Let *H*^(l-1)^ denote the matrix of sequence states entering Transformer layer *l*. For sequence-attention head *r*, queries, keys and values were obtained as

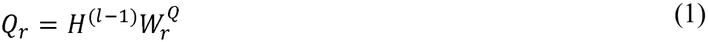

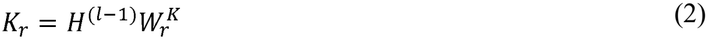

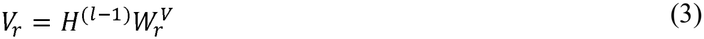

Rotary position transformations were applied to the queries and keys before scaled dot-product attention:

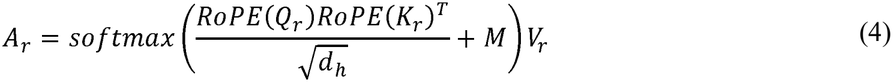

Here r ∈ {1,…,20}indexes the sequence-attention heads; 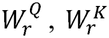 and 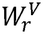are learned projections; *d*_*h*_ = 1280/20 is the dimension of each head; *M* is the additive padding mask, equal to zero at valid key positions and negative infinity at padded positions; and the softmax is taken over key positions. The superscript *T* denotes matrix transpose, and *A*_*r*_ is the context produced by head *r*. The head outputs were concatenated and passed through the attention output projection. Each block then applied post-residual layer normalization:

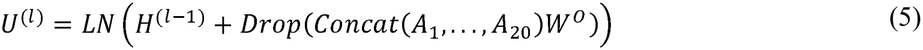

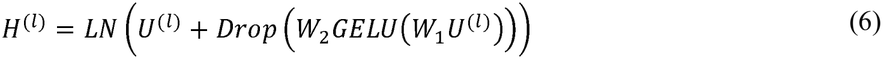

In these expressions, *W*^0^ is the multi-head output projection; *w*_1_ and *w*_2_ expand the hidden width from 1,280 to 5,120 and project it back to 1,280; *Drop* denotes dropout with probability 0.1; U^(*l*)^is the state after attention; and *H*^(*l*)^ is the output of layer *l*. The input to the first layer was the dropped-out token embedding, and a final layer normalization was applied after layer 26. In each pair-aware layer, the pair feedback Δ *h*_*i*_ defined below was added to the sequence state before the same Transformer computa-bounded tion.

#### Native retrieval of sparse partners

The Native Retriever operated at the interface between the conventional and pair-aware blocks. Given hidden states *h*_*i*_ it formed bias-free query and key vectors of dimension *d*_*p*_ = 128, normalized them to unit length, and scored a directed edge from anchor *i* to candidate partner *j* as

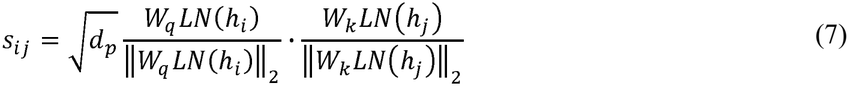

Here *sij* is the retrieval score, *w*_*q*_ and *w*_*k*_ are the learned query and key projections, respectively. Special tokens, padding and N tokens were excluded from retrieval, and candidates separated by fewer than four sequence positions were masked. The retriever selected the top *K*= 8 partners for each eligible anchor. Selection was sequence-derived; folding proposals were never inserted into the native topset. The re-sulting directed edges persisted through all eight pair-aware layers. We fixed K = 8 as a sparsity budget that retained multiple competing partners while keeping pair computation proportional to sequence length. We did not treat eight as an empirically optimal value.

#### Persistent pair-state representation

For each retrieved edge (*i,j*), RIPPLE initialized a 128-dimensional pair state. Layer-normalized end-point states were first projected into pair space, *u*_*i*_ = *W*_*u*_*LN*(*h*_*i*_), where *u*_*i*_ is the projected state of end-point *i* and *w*_u_ is the learned endpoint projection. The initialization combined symmetric endpoint fea-tures with a learned logarithmic separation embedding:

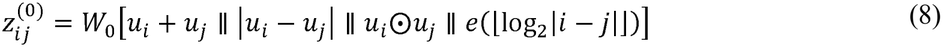

Here 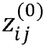 is the initial pair state, *w*_0_ is its learned output projection and *e*(*b*) is the learned embedding of separation bin *b*. The quantity | *u*_*i*_ − *u*_*j*_|is the component-wise absolute endpoint difference, whereas | *i* − *j* |is the integer sequence separation; ⌊ log_2_ | *i* −*j* |⌋ maps that separation to its logarithmic bin. At each pair-aware layer *l*, the state was updated using the current projected endpoint representations *u*_*i*_, and *u*_*j*_:

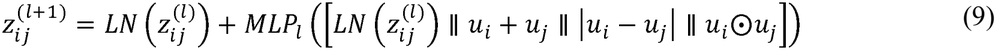

In this update, *MLP*_*l*_ is the layer-specific multilayer perceptron, and the addition to 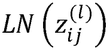is the resid-ual pair-state update that produces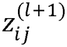. Thus the edge memory was propagated rather than recon-structed independently at every layer. The objective branch used the same pair-state initialization and update parameters for its shadow edges, but shadow states did not write to sequence hidden states.

#### Bounded pair-to-token feedback

Pair states influenced anchor representations through four-head sparse attention, with 32 dimensions per pair head. For each anchor and head, the model combined a query-key compatibility term with a learned state-dependent bias. A learned null partner was included in the softmax. If *π*_*ijm*_ is the resultingprobability for edge (*i,j*) in head *m*, the candidate weights were scaled so that their total non-null mass could not exceed *c*_*m*_ = 0.25:

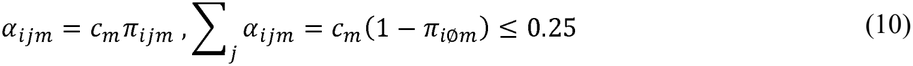

The value for an edge combined the gathered partner value with a projection of its pair state. The four head-specific contexts were concatenated and projected to the sequence width. A learned sigmoid gate, conditioned on the normalized anchor state and pair context, modulated the update. To prevent the relational branch from dominating the backbone, its update direction was normalized and its magnitude was limited relative to the anchor norm:

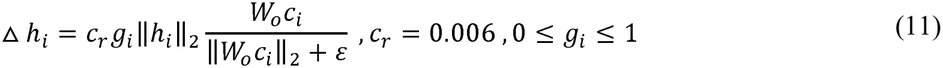

Here *c*_*i*_ is the concatenated multi-head pair context for anchor *i, W*_0_projects that context to the sequence width, *g*_*i*_ is the learned sigmoid gate and _*cr*_ is the fixed residual-magnitude cap. The positive constant *ε* prevents division by zero, and Δ *h*_*i*_ is the resulting feedback vector added to *h*_*i*_. The update was added before the corresponding Transformer block. The pair-mass cap and residual cap were fixed.

### Pairwise objective branch

The objective branch asked whether modelling two masked endpoints jointly improved their likelihood relative to predicting them independently. A linear head first produced four-way A/U/C/G marginal distributions *p*_*i*_(*a*) and *p*_*j*_(*b*)Their independent joint distribution was

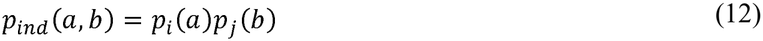

Here *a,b* ∈ {*A,U,C,G*}denote possible endpoint bases, and *p*_*ind*_ (*a,b*) is the factorized probability of the ordered pair (*a,b*).

A second head projected the final shadow pair state to 16 ordered nucleotide-pair logits. To ensure that the pair model captured coupling rather than changing the single-position predictions, the resulting coupling matrix was double-centred, clipped, and combined with the independent log probabilities. Thirty-two iterations of iterative proportional fitting ^40^ restored the two marginal distributions. For the observed endpoint bases (*y*_*i*_,*y*_*j*_), the pair loss was

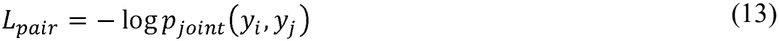

Here *y*_*i*_ and *y*_j_ are the observed masked bases, *p*_*joint*_ (*y*_*i*_,*y*_*j*_) is their fixed-marginal joint probability and *L*_*pair*_ is the pairwise negative log-likelihood. We defined pair utility as the detached reduction in negative log-likelihood:

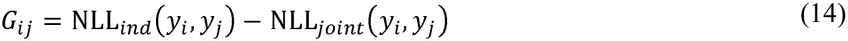

Here NLL_*ind*_ (*y*_*i*_,*y*_*j*_)and NLL_*joint*_ (*yi,yj*) are the negative log-likelihoods of the observed ordered pair under *p*_*ind*_ and *p*_*joint*_, respec0074ively, and *G*_*ij*_ is their detached difference for edge (*i,j*). Gain is a model-dependent training signal, not a calibrated base-pair probability and not a structural annotation. Positive Gain indicates that the learned coupling improved prediction for that masked example; negative Gain indicates that it worsened prediction.

### Pretraining

#### Pretraining corpus and leakage control

Pretraining used RNACentral release 26.^41^ After normalization and length filtering, the corpus contained 32,476,549 sequences before exact-overlap exclusion, leaving 32,363,130 sequences for pretraining. Records shorter than 8 nt or longer than 1,024 nt were excluded. The fold files were streamed sequentially through a shuffle buffer of 20,000 records. We retained the natural length distribution and did not sample length bins uniformly. Pretraining filters excluded exact sequence identifiers and exact nucleotide sequences appearing in the sequence-validation collection, the retrieval diagnostic sets or the downstream datasets. The filtering decision and reason were written to an audit trail. These checks protected against exact overlap.

#### Pair-guided masking and objective-edge sampling

The ordinary masked-language-modelling target was 15% of biological tokens. On 20% of batches, ordinary masked positions were sampled in contiguous spans with mean span length 4; otherwise they were sampled independently. Standard selected positions used the BERT-style 80/10/10 replacement rule:^42^ 80% were replaced with <mask>, 10% with a random canonical base and 10% were left unchanged. Both endpoints of every selected objective edge were always replaced with <mask>. Because endpoint masking took precedence, short sequences with many objective edges could exceed the ordinary 15% target, subject to an absolute endpoint-mask budget of approximately 1,500 tokens per batch.

The number of objective edges per sequence was capped at 16, 24, 32 or 40 for lengths below 256, 256-383, 384-511 or 512-1,024 nt, respectively. Anchors were selected from the sequence and shuffled without reference to predicted structure. Each anchor contributed at most two edges, each partner was used by at most two objective edges, duplicate unordered edges were removed and ∣ *i* −*j*∣ ≥4 was required.

For each eligible anchor proposal, the sampler attempted to use a partner from an RNAfold minimum-free-energy structure with probability 0.5.^43^ If no legal proposal was available, it sampled a legal partner from the sequence. Multiple bracket types in dot-bracket records were parsed. RNAfold was therefore a candidate generator, not a hard target, and the Gain objective determined whether a proposed relation was useful.

#### Pretraining objectives

RIPPLE was optimised with masked-language-modelling, pairwise and retrieval losses. The first trained contextual nucleotide representations, the second trained fixed-marginal coupling for sampled endpoint pairs and the third distilled pair utility into the sequence-only Native Retriever.

#### Masked-language-modelling loss

A separate bias-free linear head mapped the final sequence state at every position to logits over the nine-token vocabulary; its weights were not tied to the input embedding. For corrupted input *x*^*mask*^, the predicted probability of vocabulary token *ν* at position *I* was

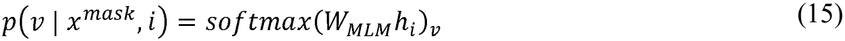

Here *w*_*MLM*_ is the language-modelling projection, *h*_i_ is the final sequence state. Let *M* contain all selected ordinary masking positions and all endpoints of sampled objective edges. The masked-language-modelling loss was the mean cross-entropy over these positions:

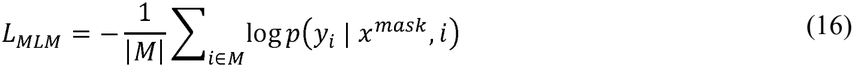

Here |*M*| is the number of supervised positions and *y*_*i*_ is the original token at position *i*. Padding and unselected positions were assigned the ignore label and did not contribute to *L*_*MLM*_ .

#### Pairwise loss

Let *E* denote the sampled objective edges in a batch. The pairwise objective averaged the per-edge fixed-marginal negative log-likelihood defined above:

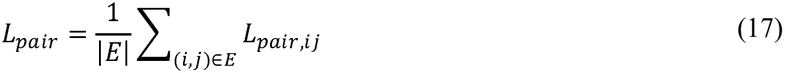

Here | *E*| is the number of objective edges. When no objective edge was present, this term was set to zero. Because both endpoint marginals came from the MLM logits and iterative proportional fitting preserved them *L*_*pair*_ trained pair-specific coupling without replacing the single-position predictions.

#### Gain-distilled retrieval loss

Objective edges sharing an anchor formed a supervision group *A*_*i*_. De-tached Gains were clipped and converted to a teacher distribution; Native Retriever scores produced the student distribution:

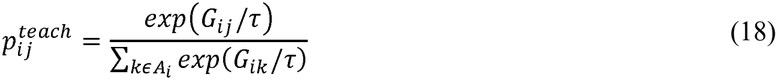

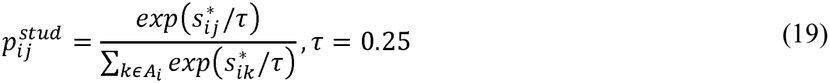

Here *G*_*ij*_ is the Gain, 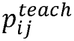is the teacher probability, 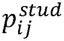 is the student probability, indexes *k* candi-date partners in the same anchor group and is the shared temperature. The score 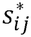 equals the Native Retriever score *s*_*ij*_, except that its gradient was stopped when edge (*i,j*) was already present in the native top-8 set. This retained existing native candidates as score references while directing gradients towards useful missed candidates.

Confidence weighting required both a missed edge above a step-dependent Gain margin mG and supporting positive evidence within the group. Let *G*_(1)_ and *G*_(2)_ denote the largest and second-largest positive missed gains. The group confidence was

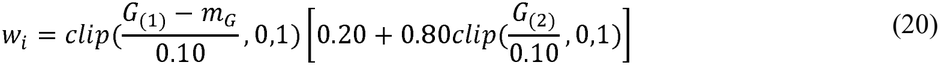

Here *W*_*i*_ is the confidence for anchor group *i*. The constant 0.1 sets the Gain normalisation scale; 0.2 is the minimum multiplicative support and 0.8 weights the additional support contributed by *G*_(2)_. Groups with fewer than two candidates or without a missed candidate were skipped. For the remaining group set *I*, the retrieval loss was

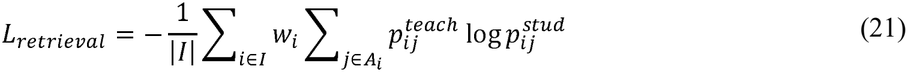

Here ∣ *I* ∣is the number of retained supervision groups. The total loss at logical training step was

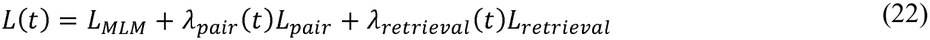

Here *λ*_*pair*_ (*t*) and *λ*_*retrieval*_ (*t*) were zero before step 5,000, increased linearly from zero to 0.1 between steps 5,000 and 25,000, and remained 0.1 thereafter. Thus, MLM had unit weight throughout training, whereas the two relational objectives were introduced progressively.

#### Pretraining optimization

RIPPLE was trained under a single schedule using fused AdamW,^44^ weight decay 0.1 and gradient-norm clipping at 0.5. The per-device batch size was 16 sequences, gradients were accumulated for two steps and training used two GPUs, giving an effective batch size of 64 sequences. The base learning rate was 2 × 10^-5^, with 2% linear warm-up followed by cosine decay over 1,500,000 steps. Pair-module parame-ters used the same base multiplier. Training used distributed data parallelism over two NVIDIA RTX 5090 GPUs (32 GB each), bfloat16 autocasting and gradient checkpointing.^45^ The host provided 50 Intel Xeon Platinum 8470Q CPU cores.

#### Retrieval diagnostics

Retrieval behaviour was monitored on BpRNA-Spot_0 and a length-filtered ArchiveII diagnostic collection. Together, these comprised 5,246 sequences and 525,504 directed reference pairs. These collections were used during development and are not described as untouched final test sets. We recorded top-1 and top-8 recall, mean reciprocal rank, area under the precision-recall curve for native and objective-forced candidates, native-entry rate, retrieval loss, gain in native membership, churn of the top-8 set and effective candidate utilisation. Diagnostic curves were used to detect collapse or instability, not to select downstream test labels.

#### Downstream evaluation

The downstream benchmark compared frozen representations under matched task heads. RIPPLE and every comparison encoder were placed in evaluation mode and executed without gradient tracking. Only the downstream head was optimised. Final-layer nucleotide states were converted to a common *L*x*d* representation, with model-specific boundary tokens removed and padding recorded separately. Sequence-level tasks used the padding-masked mean of nucleotide states. Pairwise tasks retained all nucleotide states.

The comparison encoders were RiNALMo-giga,^8^ ERNIE-RNA,^9^ RNA-FM,^12^ NucleicBERT,^13^ RNAErnie,^7^ RNA-MSM,^11^ BiRNA-BERT,^14^ RNABERT,^15^ RNAret,^16^ Uni-RNA,^46^ and HydraRNA.^47^ RNA-MSM received each RNA as a depth-one multiple-sequence alignment, so no homologous sequences were supplied. All other encoders received their native one-sequence tokenisation. All pretrained encoders were evaluated using the same benchmark sources, predefined dataset partitions and task-specific prediction head architectures. Model-specific tokenisation and supported context lengths were respected during input processing. For pairwise prediction tasks, structural annotations were adjusted to the retained sequence regions, and sequences exceeding the supported length were excluded where truncation was not applica-ble. All neural heads used a learning rate of 10-3, at most 30 epochs, and early stopping after 3 epochs without validation improvement. Dataset definitions and split assignments were fixed. Benchmark source records, processed splits and documented eligibility criteria are collated in **Supplementary Table 1**.

### ncRNA family classification

We used the 88-class ncRNA classification benchmark derived from Rfam by ncrna-deep.^48^ The dataset was constructed by retaining short ncRNAs and excluding families strongly distinguishable by sequence length or represented by fewer than 400 sequences. Sequences were partitioned within families, with sequence-similarity filtering and subsequent class balancing during dataset preparation. The resulting splits contained 105,864 training, 17,324 validation and 25,342 test sequences. All 88 families were represented in every partition, making this a within-family rather than family-held-out evaluation. Nucleotide embeddings from the frozen encoder were masked-mean-pooled, excluding special and padding tokens, and passed through a two-layer classifier with 256 hidden units, GELU activation and 88 output classes. We optimised multiclass cross-entropy using AdamW with weight decay 0.01 and retained the checkpoint with the highest validation macro-F1.

### Mean ribosome load prediction

Mean ribosome load (MRL) prediction used the massively parallel 5’ untranslated region assay of Sample *et al*. ^49^ (GEO accession GSM4084997). We implemented two frozen-encoder evaluations. The linear probe used all 106,530 assay sequences. Padding-masked mean embeddings were standardised within each training fold and fitted by ridge regression ^50^ in five shuffled outer folds. Within each outer training partition, 12.5% of examples formed an inner validation set. The regularisation coefficient was selected by inner-validation *R*^2^ from *α* ∈ {10 ^−2^,10 ^−1^,1,10,10^2^,10^3^,10^4^,10^5^,10^6^}. The reported metrics were *R*2, Pearson and Spearman correlations.

The nonlinear probe used the Optimus split of 76,319 random training sequences, 7,600 random validation sequences and 7,600 human test sequences. Each nucleotide state was projected to 32 channels and processed by six one-dimensional residual blocks.^51^ Each block contained two bias-free width-three convolutions, each followed by instance normalisation and ELU activation. Padding-masked global mean pooling, dropout of 0.2 and a linear output produced the MRL estimate. Targets were standardised using the training-set mean and standard deviation. The head was optimised by mean-squared error with Adam, and the checkpoint with the highest validation *R*2 was retained. Predictions were transformed back to the original scale and clipped at zero before evaluation.

### Secondary-structure prediction

Secondary-structure evaluation used four protocols. The SPOT-RNA/bpRNA split contained 10,814 training (TR0), 1,300 validation (VL0) and 1,305 test (TS0) sequences.^5,52^ The TS0-trained head was also evaluated without retraining on 5,401 bpRNA-new sequences, a family-wise generalisation set derived from Rfam 14.2.^53^ The local RIVAS files contained 3,166 training, 592 validation and 430 test sequences;^54^ these corresponded to TrainSetA, the available TestSetA-derived validation file and TestSetB, respectively. ArchiveII, originally distributed with the work of Sloma and Mathews,^55^ contained 3,950 sequences from ten families in our processed checkout and was evaluated by leave-one-family-out trans-fer. For each held-out family, the remaining families were shuffled with seed 42 and divided into 90% head training and 10% validation data.

For every nucleotide pair (*i,j*), the decoder concatenated [*h*_*i*_ || *h*_*j*_ ] and projected it to 64 channels. Two two-dimensional residual blocks followed. Each block comprised a 1x 1 convolution, a 3×3 convolution and a second 1×1 convolution, with instance normalisation and ReLU after every convolution. A final 3×3 convolution produced one logit per ordered pair. The strict upper triangle was mirrored to produce a symmetric map. The head was optimised with Adam, binary cross-entropy over the unpadded upper triangle and gradient-norm clipping at 1.0.

The decision threshold was selected on the validation set from 0.01 to 0.29 in increments of 0.01. Decoding excluded pairs separated by fewer than four positions and noncanonical pairs other than A-U, G-C and G-U. Candidate pairs above the threshold were greedily selected in descending probability order, with at most one partner per nucleotide. The validation criterion was mean relaxed F1, where a predicted pair was accepted within a one-nucleotide displacement of a reference pair. Scores on ArchiveII were first averaged within each held-out family and then macro-averaged across families.

### Three-dimensional contact prediction

Three-dimensional contact evaluation followed the RNAcontact benchmark,^56^ which provides 211 development and 80 test structures. We required an exact match between FASTA length and the number of residues with coordinates and applied the encoder-specific length limit. Twenty-one structures failed these checks, leaving 194 development and 76 test RNAs. A seed-42 permutation divided the retained development set into 155 training and 39 validation RNAs. PDB ATOM and HETATM records were read for each residue. Blank or A alternate locations were retained, and the highest-occupancy coordinate was selected when an atom had multiple records. A pair was labelled as a contact when its minimum interresidue atom distance was at most 8 Å.

We evaluated a strong nonlinear head and a lightweight probe. The strong head used the same 64-channel, two-block two-dimensional residual architecture as the secondary-structure decoder. The probe projected each nucleotide to 64 dimensions, formed a bilinear dot-product matrix and added a learned scalar embedding of sequence separation, clipped at 32 positions. A 3x 3 convolution smoothed the probe logits. Both outputs were symmetrised from the strict upper triangle.

Contact heads were optimised with Adam and binary cross-entropy over valid upper-triangular pairs with |*i* −*j*| ≥ 4. Gradients were clipped at norm 1.0. Checkpoint selection used the mean of the available validation metrics. Test evaluation was restricted to long-range pairs with |*i* −*j*| ≥ 24. Within each RNA, predictions were ranked, and precision was calculated at *L*. Dataset-level values were unweighted means over RNAs. Here, *P@L* denotes the fraction of true contacts among the top *L* predicted long-range nucleotide pairs for an RNA of length *L*.

### Multispecies splice-site prediction

Splice-site recognition used 400-nt windows from the multispecies Spliceator GS_1 benchmark.^57^ The original training and test partitions were used for head training and validation, respectively, without species-specific filtering. The donor splits contained 17,556 training and 4,389 validation windows, and the acceptor splits contained 17,723 training and 4,431 validation windows. External evaluation used separately prepared datasets containing 20,000 balanced windows each from Danio rerio, Drosophila melanogaster, Caenorhabditis elegans and Arabidopsis thaliana. Donor and acceptor heads were trained and evaluated separately.

The checkpoint with the highest validation F1 was retained, and a sigmoid threshold of 0.5 converted logits to labels. The trained head was then applied to each external test dataset without further adaptation.

## Data availability

The RNA sequences used for pretraining were obtained from RNAcentral release 26 (https://ftp.ebi.ac.uk/pub/databases/RNAcentral/releases/26.0/sequences/). The ncRNA family classification benchmark is available from the ncrna-deep repository (<https://github.com/bioinformaticssannio/ncrna-deep>). The mean ribosome loading data were obtained from the study by Sample et al. (2019), GEO accession GSM4084997 (https://www.ncbi.nlm.nih.gov/geo/query/acc.cgi?acc=GSM4084997). The RNA secondary-structure benchmarks include the bpRNA-1m-derived SPOT-RNA TR0/VL0/TS0 splits, bpRNA-new, RIVAS and ArchiveII-1024. The original bpRNA-1m collection is available at https://bprna.cgrb.oregonstate.edu/download.php, and the SPOT-RNA datasets are described at https://github.com/jaswindersingh2/SPOT-RNA. Corresponding processed dataset releases are hosted by MultiMolecule: bpRNA-spot-0 (https://huggingface.co/datasets/multimolecule/bprna-spot-0), bpRNA-new (https://huggingface.co/datasets/multimolecule/bprna-new), RIVAS (https://huggingface.co/datasets/multimolecule/rivas) and ArchiveII-1024 (https://huggingface.co/datasets/multimolecule/archiveii.1024). Original distributions of bpRNA-new, RIVAS and ArchiveII are also available through the MXfold2 release archive (https://github.com/mxfold/mxfold2/releases/tag/v0.1.1). The bpRNA-new benchmark was derived from Rfam release 14.2 (https://ftp.ebi.ac.uk/pub/databases/Rfam/14.2/). The RNA 3D contact prediction benchmark is available from RNAcontact (https://yanglab.qd.sdu.edu.cn/RNAcontact/benchmark/), with the underlying experimental structures deposited in the Protein Data Bank (https://www.rcsb.org/). The Spliceator GS_1 benchmark is associated with the Spliceator project, whose training and testing datasets are distributed through its official download page (https://bigest-icube.fr/spliceator/download).

## Code availability

The RIPPLE source code, environment specifications, training, evaluation configurations and final pretrained model weights are made available through the RIPPLE repository (https://github.com/ABILiLab/RIPPLE).

## Acknowledgments

F.L. is supported by the Australian National Health and Medical Research Council Investigator grants [2041439]. The South Australian immunoGENomics Cancer Institute (SAiGENCI) received grant funding from the Australian Government.

## Author information

### Contributions

F.L. conceptualised and supervised the project with help from C.S. and L.C.. H.Y. designed the model with guidance from F.L.. H.Y., G.X., and R.Z. developed the software with guidance from F.L.. All authors contributed ideas to the work and assisted in manuscript editing and revision. All authors read and approved the final manuscript.

### Corresponding authors

Fuyi Li, Christopher J Sweeney, Lachlan JM Coin

## Declarations

### Ethics approval and consent to participate

Not applicable.

### Competing interests

The authors declare no competing interests.

## Notes

### Competing Interest Statement

The authors have declared no competing interest.

