## Supplementary Material for "RIPPLE: learning nucleotide relations in an RNA foundation model"

**Supplementary Methods**

### 1. Dataset provenance and overlap control

The sources, processed releases and evaluation partitions for all downstream benchmarks are listed in Supplementary Table 1. Sample counts refer to the analyzed releases rather than to identically named original archives.

During pretraining-data preparation, 32,476,549 RNAcentral release 26 records met the 8 - 1,024-nt length criterion. Exact normalized-sequence or identifier matches to the indexed evaluation collections were excluded (113,419 records), leaving 32,363,130 sequences. This procedure addressed direct record reuse but did not establish the absence of distant homology.

### 2. Benchmark evaluation

Supplementary Tables 2-7 report 26 evaluation metrics for RIPPLE and 11 comparison encoders. Contact P@L is averaged at the RNA level, whereas ArchiveII secondary-structure scores are averaged across held-out families. Model-specific context-length limits can alter the RNAs eligible for evaluation.

### 3. Mechanistic analyses and statistics

For Fig. 2c, family-neighbourhood purity was computed in the original embedding space, while the neighbourhood-size sensitivity analysis in Fig. 2f used two-dimensional projections and a sequence-length/GC-content control. In Fig. 3, negative pairs were sampled randomly or matched to annotated pairs by sequence separation; distance matching did not additionally constrain pair chemistry (Supplementary Tables 8-13).

For the 16-state analysis in Fig. 4, nucleotide identities were changed at fixed endpoint positions; Watson-Crick versus mismatch contrasts excluded GU states. For Fig. 4f, pair separation was grouped as <32, 32-63, 64-127 and ≥128 nt. The compensatory-rescue display comprised 144 RNAs balanced across four data sources. In Fig. 5, Recall@16 is an offline ranking diagnostic rather than the deployed Top-8 candidate graph; the Fig. 6 examples are illustrative (Supplementary Tables 14-18).

Confidence intervals were estimated by RNA-level bootstrap to preserve dependence among pairs from the same RNA: 2,500 resamples for Fig. 3 and 4,000 for Figs. 4 and 5. Pair Gain and retrieval results use RNA-level summaries and AUPRC is computed at the edge level.

**Supplementary Tables**

Supplementary Table 1. Benchmark sources, processed splits and documented eligibility

Counts refer to processed dataset releases. RIVAS validation contains 592 retained TestSetA records; ArchiveII is the length-limited release. Eligibility may differ among encoders with different context limits. The original GS_1 training and test partitions were used for training and validation, respectively; no species-specific filtering was applied.

| **Benchmark** | **Source / processed release** | **Recorded split and eligibility** |
| --- | --- | --- |
| ncRNA classification | ncrna-deep; Rfam-derived 88 families | 105,864 train / 17,324 val / 25,342 test; all 88 labels in each split |
| MRL: Ridge | GEO GSM4084997; Optimus 5' UTR assay | 106,530 sequences; five shuffled outer CV folds |
| MRL: ResNet | GEO GSM4084997; random/human subsets | 76,319 random train / 7,600 random val / 7,600 human test |
| Secondary: TS0 | bpRNA-1m / SPOT-RNA; MultiMolecule bprna-spot-0 | TR0 10,814 / VL0 1,300 / TS0 1,305 |
| Secondary: bpRNA-new | MXfold2 / Rfam 14.2; MultiMolecule bprna-new | External test 5,401; the TS0-trained head is used without retraining |
| Secondary: RIVAS | MXfold2 Rivas; MultiMolecule rivas | 3,166 train / 592 val / 430 test; val retained from original 697-record TestSetA |
| Secondary: ArchiveII | ArchiveII / MXfold2; MultiMolecule archiveii.1024 | 3,950 sequences; 10 leave-one-family-out folds |
| RNA 3D contacts | RNAcontact benchmark; PDB structures | 211/80 original dev/test; 194 dev (155/39 train/val) and 76 test retained |
| Splice: donor | Spliceator GS_1, multispecies; 400-nt windows | 17,556 train / 4,389 val; four external species test sets |
| Splice: acceptor | Spliceator GS_1, multispecies; 400-nt windows | 17,723 train / 4,431 val; four external species test sets |

Supplementary Table 2. ncRNA family classification: complete 88-family benchmark metrics

Macro-F1 is the primary ncRNA endpoint in main Table 1; accuracy is shown separately. Bold indicates the best value and underlining the second-best value within each metric. Values are rounded to three decimals; rankings use the unrounded scores.

| **Model** | **Accuracy** | **Macro-F1** | **MCC** | **AUROC** |
| --- | --- | --- | --- | --- |
| RNABERT | 0.636 | 0.533 | 0.623 | 0.856 |
| RNA-FM | 0.951 | 0.840 | 0.945 | 0.999 |
| RNA-MSM | 0.713 | 0.532 | 0.697 | 0.957 |
| Uni-RNA | 0.744 | 0.573 | 0.733 | 0.985 |
| RiNALMo | 0.952 | 0.851 | 0.950 | 0.999 |
| ERNIE-RNA | 0.967 | 0.906 | 0.961 | **1.000** |
| RNAErnie | 0.896 | 0.720 | 0.889 | 0.998 |
| BiRNA-BERT | 0.867 | 0.648 | 0.858 | 0.994 |
| HydraRNA | 0.824 | 0.632 | 0.812 | 0.993 |
| NucleicBERT | 0.806 | 0.565 | 0.794 | 0.988 |
| RNAret | 0.787 | 0.595 | 0.775 | 0.990 |
| RIPPLE | **0.974** | **0.925** | **0.969** | 1.000 |

Supplementary Table 3. Mean ribosome loading: ridge-regression cross-validation

Results are from five-fold ridge regression; Spearman is the primary metric in main Table 1. Bold indicates the best value and underlining the second-best.

| **Model** | **R²** | **Pearson** | **Spearman** |
| --- | --- | --- | --- |
| RNABERT | 0.101 | 0.323 | 0.359 |
| RNA-FM | 0.177 | 0.421 | 0.423 |
| RNA-MSM | 0.114 | 0.337 | 0.333 |
| Uni-RNA | 0.191 | 0.439 | 0.466 |
| RiNALMo | 0.219 | 0.470 | 0.503 |
| ERNIE-RNA | 0.172 | 0.415 | 0.418 |
| RNAErnie | 0.186 | 0.432 | 0.436 |
| BiRNA-BERT | 0.176 | 0.420 | 0.423 |
| HydraRNA | **0.257** | **0.514** | **0.561** |
| NucleicBERT | 0.140 | 0.375 | 0.385 |
| RNAret | 0.147 | 0.383 | 0.384 |
| RIPPLE | 0.231 | 0.485 | 0.485 |

Supplementary Table 4. Mean ribosome loading: nonlinear residual-network regression

The ResNet evaluation uses the Optimus random-training and human-test split. R² is the primary metric in main Table 1; Bold indicates the best value and underlining the second-best.

| **Model** | **R²** | **Pearson** | **Spearman** |
| --- | --- | --- | --- |
| RNABERT | 0.651 | 0.828 | 0.785 |
| RNA-FM | 0.659 | 0.824 | 0.768 |
| RNA-MSM | 0.603 | 0.787 | 0.755 |
| Uni-RNA | 0.613 | 0.788 | 0.705 |
| RiNALMo | 0.557 | 0.751 | 0.678 |
| ERNIE-RNA | 0.656 | 0.813 | 0.762 |
| RNAErnie | 0.594 | 0.773 | 0.718 |
| BiRNA-BERT | 0.666 | 0.819 | 0.771 |
| HydraRNA | 0.510 | 0.735 | 0.655 |
| NucleicBERT | 0.597 | 0.777 | 0.725 |
| RNAret | 0.472 | 0.710 | 0.639 |
| RIPPLE | **0.689** | **0.831** | **0.802** |

Supplementary Table 5. RNA secondary-structure prediction: complete benchmark metrics

F1 and exact pair-matching F1 are reported separately; main Table 1 uses exact F1. ArchiveII results are averaged across held-out families. Bold indicates the best value and underlining the second-best.

| **Model** | **TS0 F1** | **TS0 exact F1** | **bpRNA-new F1** | **bpRNA-new exact F1** | **ArchiveII F1** | **ArchiveII exact F1** | **RIVAS F1** | **RIVAS exact F1** |
| --- | --- | --- | --- | --- | --- | --- | --- | --- |
| RNABERT | 0.434 | 0.404 | 0.408 | 0.399 | 0.279 | 0.272 | 0.384 | 0.369 |
| RNA-FM | 0.620 | 0.579 | 0.447 | 0.436 | 0.296 | 0.290 | 0.521 | 0.506 |
| RNA-MSM | 0.475 | 0.442 | 0.420 | 0.410 | 0.267 | 0.261 | 0.367 | 0.348 |
| Uni-RNA | 0.532 | 0.497 | 0.480 | 0.470 | 0.362 | 0.353 | 0.442 | 0.424 |
| RiNALMo | 0.638 | 0.595 | 0.470 | 0.459 | 0.601 | 0.590 | 0.610 | 0.588 |
| ERNIE-RNA | 0.690 | 0.660 | **0.570** | **0.559** | 0.589 | 0.576 | 0.694 | 0.655 |
| RNAErnie | 0.538 | 0.502 | 0.424 | 0.413 | 0.402 | 0.395 | 0.424 | 0.408 |
| BiRNA-BERT | 0.522 | 0.490 | 0.464 | 0.456 | 0.306 | 0.299 | 0.461 | 0.442 |
| HydraRNA | 0.557 | 0.521 | 0.489 | 0.479 | 0.372 | 0.364 | 0.451 | 0.433 |
| NucleicBERT | 0.600 | 0.566 | 0.448 | 0.439 | 0.465 | 0.458 | 0.487 | 0.474 |
| RNAret | 0.422 | 0.394 | 0.403 | 0.394 | 0.341 | 0.334 | 0.363 | 0.348 |
| RIPPLE | **0.718** | **0.671** | 0.522 | 0.509 | **0.610** | **0.599** | **0.697** | **0.676** |

Supplementary Table 6. Intramolecular RNA 3D contacts: strong decoder and lightweight probe

Contact metrics are evaluated on long-range intramolecular pairs. Strong P@L and Probe P@L appear in main Table 1. Bold indicates the best value and underlining the second-best.

| **Model** | **Strong P@L** | **Strong P@L/2** | **Strong AUROC** | **Probe P@L** |
| --- | --- | --- | --- | --- |
| RNABERT | 0.389 | 0.521 | 0.767 | 0.204 |
| RNA-FM | 0.518 | 0.583 | 0.794 | 0.364 |
| RNA-MSM | 0.396 | 0.522 | 0.774 | 0.275 |
| Uni-RNA | 0.459 | 0.524 | 0.787 | 0.321 |
| RiNALMo | 0.558 | 0.663 | 0.819 | 0.423 |
| ERNIE-RNA | 0.565 | **0.674** | 0.830 | 0.334 |
| RNAErnie | 0.446 | 0.528 | 0.774 | 0.325 |
| BiRNA-BERT | 0.458 | 0.581 | 0.791 | 0.191 |
| HydraRNA | 0.517 | 0.599 | 0.805 | **0.460** |
| NucleicBERT | 0.355 | 0.441 | 0.752 | 0.286 |
| RNAret | 0.379 | 0.460 | 0.755 | 0.296 |
| RIPPLE | **0.575** | 0.652 | **0.836** | 0.422 |

Supplementary Table 7. Multispecies splice-site recognition: donor and acceptor performance

Donor and acceptor results summarize prediction on four external species. Bold indicates the best value and underlining the second-best for each metric.

| **Model** | **Donor F1** | **Donor AUROC** | **Acceptor F1** | **Acceptor AUROC** |
| --- | --- | --- | --- | --- |
| RNABERT | 0.551 | 0.510 | 0.549 | 0.524 |
| RNA-FM | 0.660 | 0.628 | 0.669 | 0.658 |
| RNA-MSM | 0.631 | 0.583 | 0.725 | 0.792 |
| Uni-RNA | 0.662 | 0.647 | 0.692 | 0.686 |
| RiNALMo | 0.694 | 0.715 | 0.715 | 0.783 |
| ERNIE-RNA | 0.682 | 0.697 | 0.701 | 0.761 |
| RNAErnie | 0.660 | 0.658 | 0.545 | 0.752 |
| BiRNA-BERT | 0.647 | 0.634 | 0.677 | 0.691 |
| HydraRNA | 0.681 | 0.723 | **0.765** | 0.800 |
| NucleicBERT | 0.670 | **0.728** | 0.754 | **0.834** |
| RNAret | 0.652 | 0.612 | 0.645 | 0.665 |
| RIPPLE | **0.710** | 0.717 | 0.736 | 0.809 |

Supplementary Table 8. Family-resolved sample counts and high-dimensional 10-NN purity (main Fig. 2b,c)

High-dimensional ten-nearest-neighbour purity is calculated on the 4,766-RNA balanced subset; the counts of test RNAs refer to all 18,347 sequences. The unweighted mean family purity is 0.990.

| **Rfam family** | **Test RNAs** | **Balanced RNAs** | **10-NN purity** |
| --- | --- | --- | --- |
| RF00001 | 4,179 | 400 | 0.992 |
| RF00004 | 2,913 | 400 | 0.997 |
| RF00005 | 2,524 | 400 | 0.985 |
| RF02271 | 2,242 | 400 | 1.000 |
| RF00059 | 2,216 | 400 | 0.992 |
| RF00504 | 1,102 | 400 | 0.967 |
| RF01705 | 798 | 400 | 0.992 |
| RF00050 | 589 | 400 | 0.997 |
| RF00174 | 517 | 400 | 0.990 |
| RF00379 | 487 | 400 | 0.988 |
| RF00097 | 414 | 400 | 0.997 |
| RF00168 | 366 | 366 | 0.984 |

Supplementary Table 9. Pair Gain by annotated pair chemistry (main Fig. 3b,i)

Gain means and confidence intervals are RNA-level summaries; the positive-Gain fraction is calculated over annotated edges.

| **Pair type** | **RNAs** | **Mean Gain** | **95% CI** | **Edges** | **Gain > 0** |
| --- | --- | --- | --- | --- | --- |
| AU | 6,806 | 0.911 | 0.900 to 0.923 | 16,013 | 99.81% |
| GC | 7,834 | 0.824 | 0.815 to 0.833 | 27,027 | 99.96% |
| GU | 3,582 | -0.249 | -0.267 to -0.230 | 4,964 | 31.18% |
| other | 623 | -1.101 | -1.227 to -0.977 | 980 | 13.88% |

Supplementary Table 10. Annotated Pair Gain by structural context (main Fig. 3c)

An RNA may contribute to more than one structural context. Gain estimates are RNA-level summaries.

| **Structural context** | **RNAs** | **Mean Gain** | **95% CI** |
| --- | --- | --- | --- |
| stem | 8,026 | 0.727 | 0.717 to 0.736 |
| hairpin | 3,715 | 0.696 | 0.674 to 0.718 |
| internal | 3,830 | 0.508 | 0.490 to 0.527 |
| junction | 1,451 | 0.487 | 0.452 to 0.524 |
| pseudoknot | 443 | 0.381 | 0.323 to 0.442 |

Supplementary Table 11. Distance-stratified Gain and edge-level AUPRC (main Fig. 3d)

Gain means and confidence intervals are RNA-level estimates; AUPRC is calculated at the edge level. Distance bins follow main Fig. 3d.

| **Separation** | **RNAs** | **Edges** | **Mean Gain** | **95% CI** | **AUPRC** |
| --- | --- | --- | --- | --- | --- |
| <32 | 7,906 | 68,545 | 0.733 | 0.724 to 0.743 | 0.762 |
| 32-128 | 6,863 | 63,141 | 0.653 | 0.642 to 0.664 | 0.777 |
| 128-256 | 1,176 | 10,633 | 0.394 | 0.364 to 0.425 | 0.691 |
| >256 | 595 | 4,379 | 0.264 | 0.227 to 0.301 | 0.659 |

Supplementary Table 12. Source-stratified Pair Gain gap and native MRR (main Fig. 3h)

Gain gap is the within-RNA annotated-minus-distance-matched mean Pair Gain. Confidence intervals are calculated by RNA-level resampling.

| **Source** | **RNAs** | **Gain gap** | **95% CI, gap** | **MRR** | **95% CI, MRR** |
| --- | --- | --- | --- | --- | --- |
| BpRNA-New | 5,235 | 0.538 | 0.531 to 0.544 | 0.325 | 0.321 to 0.329 |
| BpRNA-TS0 | 1,173 | 0.302 | 0.278 to 0.327 | 0.352 | 0.341 to 0.363 |
| ArchiveII1024 | 1,249 | 0.383 | 0.365 to 0.402 | 0.310 | 0.299 to 0.319 |
| RIVAS-TS | 384 | 0.242 | 0.202 to 0.281 | 0.248 | 0.239 to 0.258 |

Supplementary Table 13. Gain-gap deciles and median native-partner MRR (main Fig. 3j)

Deciles are defined using RNA-level Gain gaps (n = 8,041). The overall correlation with native MRR is Spearman rho = 0.306 (two-sided P = 3.080 × 10^-174^).

| **Gain-gap decile** | **RNAs** | **Median Gain gap** | **Median native MRR** |
| --- | --- | --- | --- |
| 1 | 805 | -0.008 | 0.240 |
| 2 | 804 | 0.160 | 0.238 |
| 3 | 804 | 0.272 | 0.251 |
| 4 | 804 | 0.366 | 0.252 |
| 5 | 804 | 0.441 | 0.265 |
| 6 | 804 | 0.516 | 0.295 |
| 7 | 804 | 0.593 | 0.317 |
| 8 | 804 | 0.673 | 0.341 |
| 9 | 804 | 0.768 | 0.372 |
| 10 | 804 | 0.934 | 0.398 |

Supplementary Table 14. Pair Gain for all 16 ordered nucleotide states (main Fig. 4a)

All 16 ordered nucleotide combinations were evaluated at the same pair of positions within each of 6,445 RNAs. Estimates are medians across RNAs with RNA-level confidence intervals.

| **Pair state** | **Class** | **Median Gain (nats)** | **95% CI** |
| --- | --- | --- | --- |
| AU | Watson-Crick | 1.052 | 1.040 to 1.066 |
| CG | Watson-Crick | 1.038 | 1.025 to 1.050 |
| GC | Watson-Crick | 1.036 | 1.026 to 1.047 |
| UA | Watson-Crick | 1.060 | 1.045 to 1.073 |
| GU | GU wobble | -0.133 | -0.142 to -0.122 |
| UG | GU wobble | -0.205 | -0.219 to -0.194 |
| AA | mismatch | -0.572 | -0.592 to -0.541 |
| AC | mismatch | -1.048 | -1.079 to -1.022 |
| AG | mismatch | -1.516 | -1.557 to -1.479 |
| CA | mismatch | -1.029 | -1.054 to -1.007 |
| CC | mismatch | -1.116 | -1.154 to -1.079 |
| CU | mismatch | -1.581 | -1.631 to -1.534 |
| GA | mismatch | -1.485 | -1.532 to -1.436 |
| GG | mismatch | -2.281 | -2.343 to -2.219 |
| UC | mismatch | -1.641 | -1.696 to -1.596 |
| UU | mismatch | -2.120 | -2.197 to -2.062 |

Supplementary Table 15. Counterfactual compatibility contrasts and rescue by RNA source (main Fig. 4b,c,g,h)

Watson-Crick-minus-mismatch contrasts exclude GU states. The rescue subset comprises 144 RNAs balanced across four sources (36 per source).

| **Source** | **Full cohort RNAs** | **Median WC minus mismatch** | **95% CI** | **Rescue RNAs** | **Gain rescued** |
| --- | --- | --- | --- | --- | --- |
| All sources | 6,445 | 2.895 | 2.854 to 2.938 | 144 | 99.3% |
| ArchiveII | 586 | 5.343 | 5.096 to 5.588 | 36 | 100% |
| BpRNA-New | 4,425 | 2.690 | 2.656 to 2.723 | 36 | 100% |
| BpRNA-TS0 | 1,058 | 3.230 | 3.121 to 3.335 | 36 | 97.2% |
| RIVAS-TS | 376 | 3.834 | 3.587 to 3.983 | 36 | 100% |

Supplementary Table 16. Native Retriever Recall@8 across sources and helix topology (main Fig. 5b,d,e,g,h)

Recall@8 is averaged across RNAs. The analysis strata are not mutually exclusive, and their sample counts are not additive.

| **Analysis stratum** | **Category** | **RNAs** | **Mean Recall@8** | **95% CI** |
| --- | --- | --- | --- | --- |
| RNA source | ArchiveII | 586 | 0.778 | 0.763 to 0.792 |
| RNA source | BpRNA-New | 4,425 | 0.666 | 0.660 to 0.671 |
| RNA source | BpRNA-TS0 | 1,030 | 0.675 | 0.660 to 0.689 |
| RNA source | RIVAS-TS | 376 | 0.552 | 0.536 to 0.569 |
| RNA source | All sources | 6,417 | 0.671 | 0.666 to 0.675 |
| Helix position | stem interior | 6,415 | 0.791 | 0.786 to 0.796 |
| Helix position | stem edge | 6,417 | 0.533 | 0.527 to 0.539 |
| Helix position | isolated | 2,379 | 0.273 | 0.256 to 0.290 |
| Stem length (pairs) | 1 | 2,379 | 0.273 | 0.256 to 0.290 |
| Stem length (pairs) | 2 | 2,835 | 0.404 | 0.391 to 0.418 |
| Stem length (pairs) | 3-4 | 5,375 | 0.614 | 0.607 to 0.622 |
| Stem length (pairs) | ≥5 | 6,102 | 0.724 | 0.718 to 0.729 |
| Pair separation (nt) | <32 | 6,316 | 0.702 | 0.697 to 0.707 |
| Pair separation (nt) | 32-63 | 5,172 | 0.537 | 0.527 to 0.547 |
| Pair separation (nt) | 64-127 | 3,248 | 0.622 | 0.608 to 0.635 |
| Pair separation (nt) | ≥128 | 417 | 0.409 | 0.377 to 0.441 |
| Stack depth | 1 | 6,417 | 0.521 | 0.515 to 0.527 |
| Stack depth | 2 | 6,415 | 0.758 | 0.753 to 0.763 |
| Stack depth | 3 | 6,102 | 0.831 | 0.824 to 0.837 |
| Stack depth | 4+ | 4,140 | 0.861 | 0.853 to 0.870 |

Supplementary Table 17. Case-level performance and annotated-partner retrieval (main Fig. 6a-g)

Gain gap compares annotated with distance-matched false pairs; exact F1 measures the downstream secondary-structure prediction, whereas Recall@8 describes Native Retriever coverage.

| **RNA case** | **Dataset** | **Length (nt)** | **Annotated pairs** | **Exact F1** | **Gain gap** | **Recall@8** |
| --- | --- | --- | --- | --- | --- | --- |
| bpRNA_RFAM_6139 | BpRNA-TS0 | 117 | 21 | 0.755 | 1.902 | 0.952 |
| URS0000D69221_12908_1 | BpRNA-New | 88 | 19 | 0.923 | 0.925 | 0.632 |
| bpRNA_RFAM_21434 | BpRNA-TS0 | 81 | 27 | 0.941 | 1.108 | 0.963 |

Supplementary Table 18. Chemistry-resolved mean Pair Gain in illustrative RNAs (main Fig. 6h)

Values are mean Pair Gain in nats, with annotated-pair counts in parentheses. A dash indicates that no pair of the corresponding type was annotated.

| **RNA case** | **AU** | **GC** | **GU** | **Other** |
| --- | --- | --- | --- | --- |
| long-range helix | 1.258 (n=8) | 0.052 (n=10) | -0.222 (n=2) | -0.275 (n=1) |
| mixed-pair helix | 1.035 (n=6) | 0.895 (n=11) | -0.207 (n=2) | - |
| compact stem network | 1.223 (n=10) | 1.227 (n=16) | - | -3.589 (n=1) |
